# Domain model explains propagation dynamics and stability of K27 and K36 methylation landscapes

**DOI:** 10.1101/729905

**Authors:** C. Alabert, C. Loos, M. Voelker-Albert, S. Graziano, I. Forné, N. Reveron-Gomez, L. Schuh, J. Hasenauer, C. Marr, A. Imhof, A. Groth

## Abstract

Chromatin states must be maintained during cell proliferation to uphold cellular identity and genome integrity. Inheritance of histone modifications is central in this process. However, the histone modification landscape is challenged by incorporation of new unmodified histones during each cell cycle and the principles governing heritability remain unclear. Here, we take a quantitative computational modeling approach to describes propagation of K27 and K36 methylation states. We measure combinatorial K27 and K36 methylation patterns by quantitative mass spectrometry on subsequent generations of histones. Using model comparison, we reject active global demethylation and invoke the existence of domains defined by distinct methylation endpoints. We find that K27me3 on pre-existing histones stimulates the rate of *de novo* K27me3 establishment, supporting a read-write mechanism in timely chromatin restoration. Finally, we provide a detailed, quantitative picture of the mutual antagonism between K27 and K37 methylation, and propose that it stabilizes epigenetic states across cell division.

## INTRODUCTION

Most cells in a multicellular organism can be functionally very distinct despite the fact that they share the same genomic information. Cellular specialization during development is based on the ability to establish, maintain, and execute different gene expression programs. How transcriptional programs are established during development and maintained in cycling cells is a fundamental question in biology. Chromatin organization plays a fundamental role in this process, but it remains unclear how specific chromatin states are stably inherited from a mother cell to its daughters. Histone post-translational modifications are highly cell type specific and constitute an important level of chromatin regulation that controls transcription programs. Histone modifications can be inherited from mother cells through DNA replication and mitosis to daughter cells, but the mechanisms underlying this inheritance are still not fully understood.

During DNA replication, old histones H3-H4 are evicted from the parental strand and re-deposited onto the two daughter DNA strands. Histone acetylation and methylation marks are maintained on the old histones during this recycling process (Alabert et al., 2015; Scharf et al., 2009; Xu et al., 2012). Recycling is accurate and almost symmetric, such that positional information is maintained and the two strands receive close to equal contributions of old modified histones (Petryk et al., 2018; Reveron-Gomez et al., 2018; Yu et al., 2018). In parallel, newly synthesized histones are deposited to maintain nucleosome density. Because new histones are largely unmodified, except from transient di-acetylation on histone H4, histone methylation levels are diluted two-fold on the daughter strands as compared to parental chromatin. In order to maintain chromatin states across cell division, new histones become modified to resemble their neighboring old histones. This process of chromatin restoration is highly heterogeneous, taking place across the full cell cycle with modification and locus specific kinetics (Alabert et al., 2015; Reveron-Gomez et al., 2018).

Despite major advances in understanding histone dynamics coupled to DNA replication, the picture remains rudimentary with respect to the mechanisms that govern restoration and thus heritability of modifications. This partly reflects the complexity in the regulation of histone modifications, involving positive and negative cross-talk among modifications themselves and with other chromatin features (e.g. DNA methylation and sequence elements) and processes (e.g. transcription). A favored paradigm for inheritance of the repressive modifications, H3K9me3 and H3K27me3, is self-propagation through a read-write mechanism where the enzymes are activated by the presence of the modification on nearby nucleosomes (Reinberg and Vales, 2018). Structural and biochemical evidence strongly support a read-write mechanism in H3K9me3 and H3K27me3 establishment and spreading (Allshire and Madhani, 2018; Reinberg and Vales, 2018): EZH2, the catalytic subunit of PRC2 that mediates mono-, di- and tri-methylation of H3K27 (from here on denoted as K27me1/2/3) is allosterically activated by binding of pre-existing K27me3 to an aromatic cage in the EED subunit (Reinberg and Vales, 2018). This can work in trans between neighboring nucleosomes (Poepsel et al., 2018) and could allow K27me3 on recycled old histones to instruct establishment of K27me3 on new histones after DNA replication (Reinberg and Vales, 2018). Consistent with this, EED cage mutations that abrogate allosteric activation reduce K27me3 levels and spreading (Lee et al., 2018; Oksuz et al., 2018). Intriguingly, PRC2 also methylates its binding partner JARID2 on a K27 mimicking peptide that also can activate the enzyme allosterically (Sanulli et al., 2015). However, other factors including genomic features, bound RNA and H2A ubiquitination by PRC1 are likely to also contribute to K27me3 maintenance (King et al., 2018; Laugesen et al., 2019; Yu et al., 2019). In Drosophila, self-propagation is not sufficient for K27me3 maintenance as sequence elements directing PRC recruitment are also required (Coleman and Struhl, 2017; Laprell et al., 2017). In mammalian cells, CpG richness is important for PRC2 recruitment but a distinct recognition element remains to be identified (Laugesen et al., 2019). However, reintroduction of PRC2 into ESCs devoid of K27 methylation fully restore K27me3 landscape (Hojfeldt et al., 2018; Oksuz et al., 2018). This argues against a critical role for allosteric activation in establishment, but its importance in K27me3 maintenance remains debated.

K27me shows an intriguing interplay with methylation of K36, located in close vicinity on the H3 tail. H3K36 mono-, di- and tri-methylation (from here on called K36me1/2/3) occupy distinct regions of the genome much like K27me1/2/3, but is linked to transcription rather than repression. K36me2 is imposed by NSD1-3 and ASH1/1L and broadly distributed across the genome including genic and intergenic regions. K36me3 is imposed by a single enzyme, SETD2, over gene bodies and promotes transcription fidelity by restoring non-permissive chromatin state following RNA Pol II passage (Huang and Zhu, 2018). Consistent with their opposite roles in transcription, K36me3 and K27me3 are mutually exclusively distributed along chromosomes (Ernst and Kellis, 2010; Streubel et al., 2018). Intriguingly, de-regulation of these modifications can give rise to similar clinical outcomes, as Weaver syndromes (missense mutations in PRC2 subunits) and Sotos syndromes (loss-of-function mutations or deletions in NSD1 or SETD2) largely phenocopy each other (Tatton-Brown and Rahman, 2013). Both K27 and K36 are also the target of recurrent mutations in histones genes, leading to expression of so-called oncohistones (K27M and K36M) that are drivers of distinct types of pediatric cancers (Mohammad and Helin, 2017). Notably, K36 and K27 methylation landscapes are highly inter-dependent; when one modification is impaired, the other one increases and spreads (Lu et al., 2016; Oksuz et al., 2018; Stafford et al., 2018; Streubel et al., 2018). Consistent with this, a bi-directional antagonism was predicted based on computational modelling of *in vivo* K27 and K36 methylation dynamics measured by mass spectrometry in cancer cells (Zheng et al., 2012). Biochemical analysis show that K36me2/3 directly blocks PRC2 mediated K27me2/3 on the same histone tail (Schmitges et al., 2011; Yuan et al., 2011), and deposition of K36me2 was proposed to directly limit K27me3 spreading (Streubel et al., 2018). Yet, genome-wide analysis shows substantial overlap between K27me2 and K36me2 and moderate correlation between K27me3 and K36me2 (Streubel et al., 2018). Also, it remains unclear how K27me2/3 might counteract K36me2/3. Thus, the nature of the antagonistic relationship and how they may influence each other during maintenance and establishment remains puzzling.

In this study we take a quantitative approach to understand how histone modification are inherited across cell division, monitoring *in vivo* dynamics of the tightly linked histone modifications K36me and K27me. We analyze 15 distinct K27 and K36 methylation states on three different generations of histones. Using a mechanistic model for histone methylation, parameter inference and quantitative hypothesis testing, we find that domains with distinct end-point methylation states have to be introduced to fit our data. We provide a detailed, quantitative picture of K27me/K36me antagonistic relationship and reveal that co-occupancy of these modifications can be explained by their distinct establishment kinetics. We also demonstrate, using an EZH2 inhibitor, that the rate of de novo K27me3 on newly deposited histones is enhanced by pre-existing K27me3, lending strong support to the read-write model.

## RESULTS

### Slow K27me3 establishment in mouse embryonic stem cells (mESCs)

To identify the principles for how K27me and K36me patterns are propagated in mESCs, we chose a quantitative approach to measure how methylations developed over time on subsequent generations of histones (Fig. 1A). We used triple SILAC labelling with light (R0K0), medium (R6K4) and heavy (R10K8) amino acids to quantitate methylation patterns over time (Fig. 1B, S1A). This scheme allows us to measure methylation levels as they develop over time on pre-existing histones (generation 1), new histones incorporated in a narrow time-window and allowed to age (generation 2), and new histones continuously incorporated over time to reach steady state levels (generation 3) (Fig. 1B). Before switching culture medium (Fig. 1B, t = −3 hr), generation 1 histones constitute the total pool of nucleosomal histones and thus provide the total methylation level at steady state, including fully modified histones and histones that still undergo modification. We focused our analysis on combinatorial K27 and K36 methylation patterns and for comparison included analysis of H3K9 and H4K20 methylation. In steady state, the most abundant modification is K27me2 present on 34±5% (mean±std., n=6 experiments) of histone H3, while 20±2% carry K27me1 and 21±1% carry K27me3 (Fig. 1C, Fig. S1B). About 50% of histones carry K36 methylation with K36me1 on 20±4%, K36me2 on 12±2% and K36me3 on 14±4%. This is overall in good accordance with previous mass spectrometry analysis of H3 methylation in mESCs growing in 2i medium (Lee et al., 2018; Oksuz et al., 2018).

**Figure 1.**
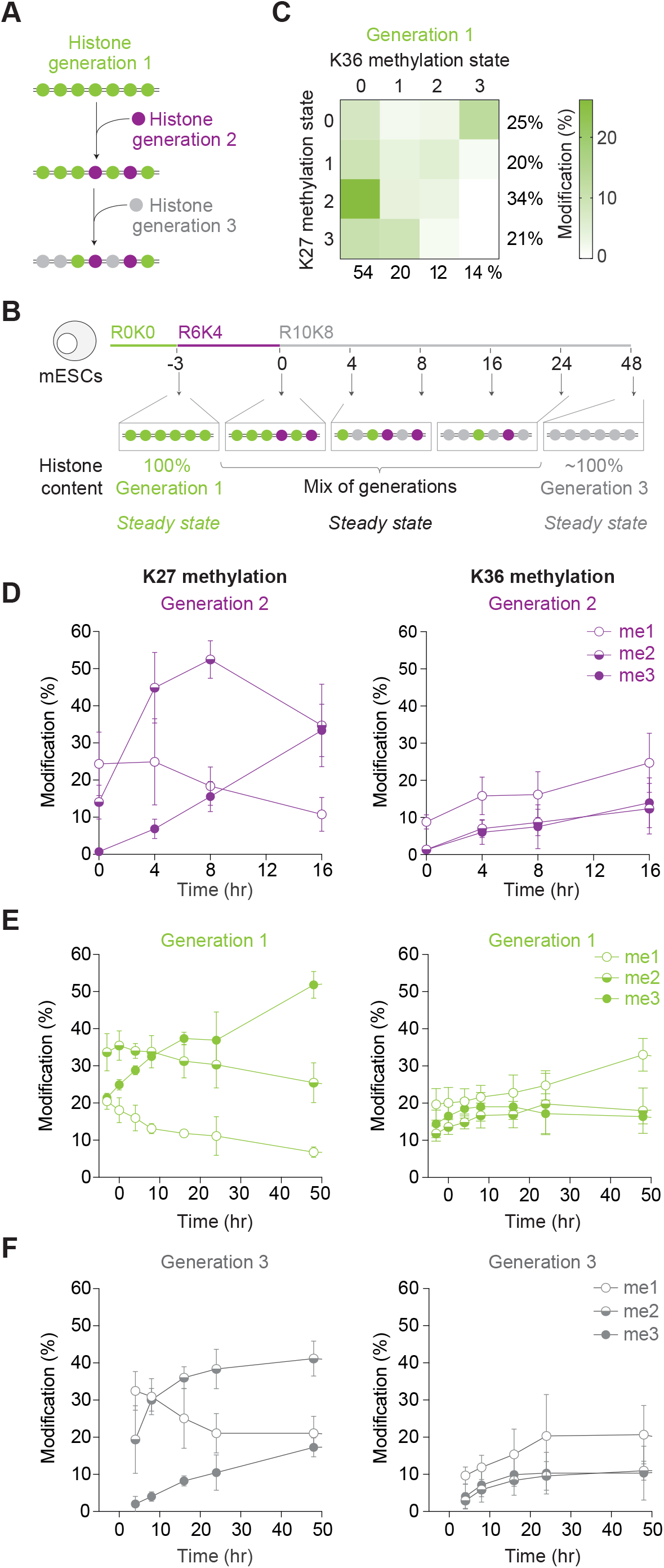
Propagation dynamics of K27 and K36 methylation in mESCs. **A**. Strategy to analyze three successive histone generations by triple SILAC. **B.** Experimental design of pulse chase SILAC labelling in asynchronous mESCs. **C.** Heatmap showing the distribution of K27 and K36 methylation states in steady-state prior to labelling. The sum of each methylation is shown on the right for K27 and at the bottom for K36. **D.** Methylation levels on generation 2 histones for K27 and K36. (me1) mono-methyl; (me2) di-methyl; (me3) tri-methyl. **E.** Methylation levels on generation 3 histones for K27 and K36. **F.** Methylation levels on generation 1 histones for K27 and K36. See also Figure S1.

We took advantage of the triple SILAC set-up to determine how K27 methylation is established on newly synthesized histones in generation 2. We observed a stepwise acquisition of K27 methylation (Fig. 1D, S1C). K27me1 occurred almost concomitant with histone incorporation and reached its maximum at the 0-hour timepoint. This was followed by a surge of K27me2 within the first 3 hours and a peak 8 hours after incorporation. K27me3 establishment occurred with a substantial delay and continued to rise for more than 16 hours after incorporation (Fig. 1D). Consistent with the latter, the analysis of generation 1 histones demonstrated that old histones continued to gain K27me3 and progressed towards a higher methylation state (Fig. 1E). As expected, generation 3 histones followed a trajectory towards the steady state levels (Fig. 1F), reaching methylation levels comparable to those of generation 1 at the beginning of the experiment (Fig. S1B). Collectively, this mirrors the pattern of K27 methylation establishment in HeLa cells (Alabert et al., 2015), although the kinetics are faster in mESCs, matching their short cell cycle of around 15 hours (Fig. S1A). The continuous methylation on old histones is specific to K27, as neither K36me3 nor H3K9me3 continues to rise on old histones beyond one cell cycle (Fig. 1F, S1E). This may reflect an inherent feature of the EZH2 enzyme that has a slow methylation rate for K27me2 to K27me3 (Justin et al., 2016; McCabe et al., 2012) and that the enzyme is challenged by the large substrate pool (in steady-state almost 75% of nucleosomal histones carry either K27me1/2/3) (Fig. 1F, Fig. S1B). mESCs contain high concentration of PRC2 (Stafford et al., 2018) probably explaining how they maintain high levels of K27me3 level through rounds of rapid cell divisions despite slow K27 tri-methylation establishment (Fig. S1B).

K36me1/2/3 establishment was continuous and slow, lasting from the time of incorporation and up to 16 hours, with K36me1 initiating shortly after incorporation and K36me2/3 delayed for a couple of hours (Fig. 1D). K36me1 and K36me2 establishment were both delayed compared to K27me1 and K27me2, respectively. These kinetics largely agrees with previous study (Zheng et al., 2012). We observed a similar acquisition pattern for K9me1/2/3 (Fig. S1E), while H4K20me1/2 show stepwise acquisition similar to K27me1/2/3 (Fig. S1F). It is not clear what governs this behavior, but it is notable that both K27me and H4K20me are highly abundant marks in mESCs 2i and that K9me show stepwise acquisition in HeLa cells where K9me2/3 levels are much higher than in mESCs 2i (Alabert et al., 2015). Curiously, the K36me2 and K36me3 kinetics were highly similar suggesting that K36me2 could be rate limiting for K36me3 establishment, in contrast to K27me where tri-methylation seems to be the slowest reaction.

### Computational model for propagation of K27 and K36 methylation

Our approach allows us to measure all combinatorial methylation states of K27 and K36 (Fig. 1C) apart from K27me3K36me3 that is below our detection limit. To understand the dynamics in the three histone generations, we employed mechanistic modelling. Such a model can describe the temporal evolution of the methylation levels. By fitting mechanistic models to our comprehensive data, we could test hypotheses on global demethylation (Reveron-Gomez et al., 2018; Zheng et al., 2012) and the previously reported negative interaction between methylations on the two residues (Schmitges et al., 2011; Stafford et al., 2018; Voigt et al., 2012; Yuan et al., 2011). For a mathematical description, we used a system of ordinary differential equations and assume that transitions between modifications follow mass action kinetics (see Supplementary Information, and Fig. S2A-B for more details on the modelling). Moreover, newly incorporated histones during cell division are assumed to be unmodified on K27 and K36 (Alabert et al., 2015; Jasencakova et al., 2010; Loyola et al., 2006). Based on these assumptions, we considered two different model variants: In the ‘global model’ (similar to the approach of Zheng et al. 2012, see Supplementary Information for a detailed description and comparison), each state develops in time due to methylation and demethylation (Fig. 2A). Since the existence of global demethylation of histones has been questioned recently (Reveron-Gomez et al., 2018), and the global model without demethylation does not fit the data (Fig. S2C), we also considered an alternative model. This model does not include demethylation and assumes that histones in certain domains can only be methylated up to a defined final state (Fig. 2B). We refer to this model as the ‘domain model’. Domains may form due to constraints in the activity of enzymes, reflecting that they are recruited and activated at specific regions of the genome (Ferrari et al., 2014; Lee et al., 2018) and leading to inhomogeneous distributions of enzymes in the nucleus. Indeed, we found that a domain model with 32 parameters and no demethylation (Fig. 2B, Supplementary Information) could explain modification dynamics in all generations as well as the global model with 30 parameters and active demethylation (Fig. 2C and Fig. S2C-E, Supplementary Information).

**Figure 2.**
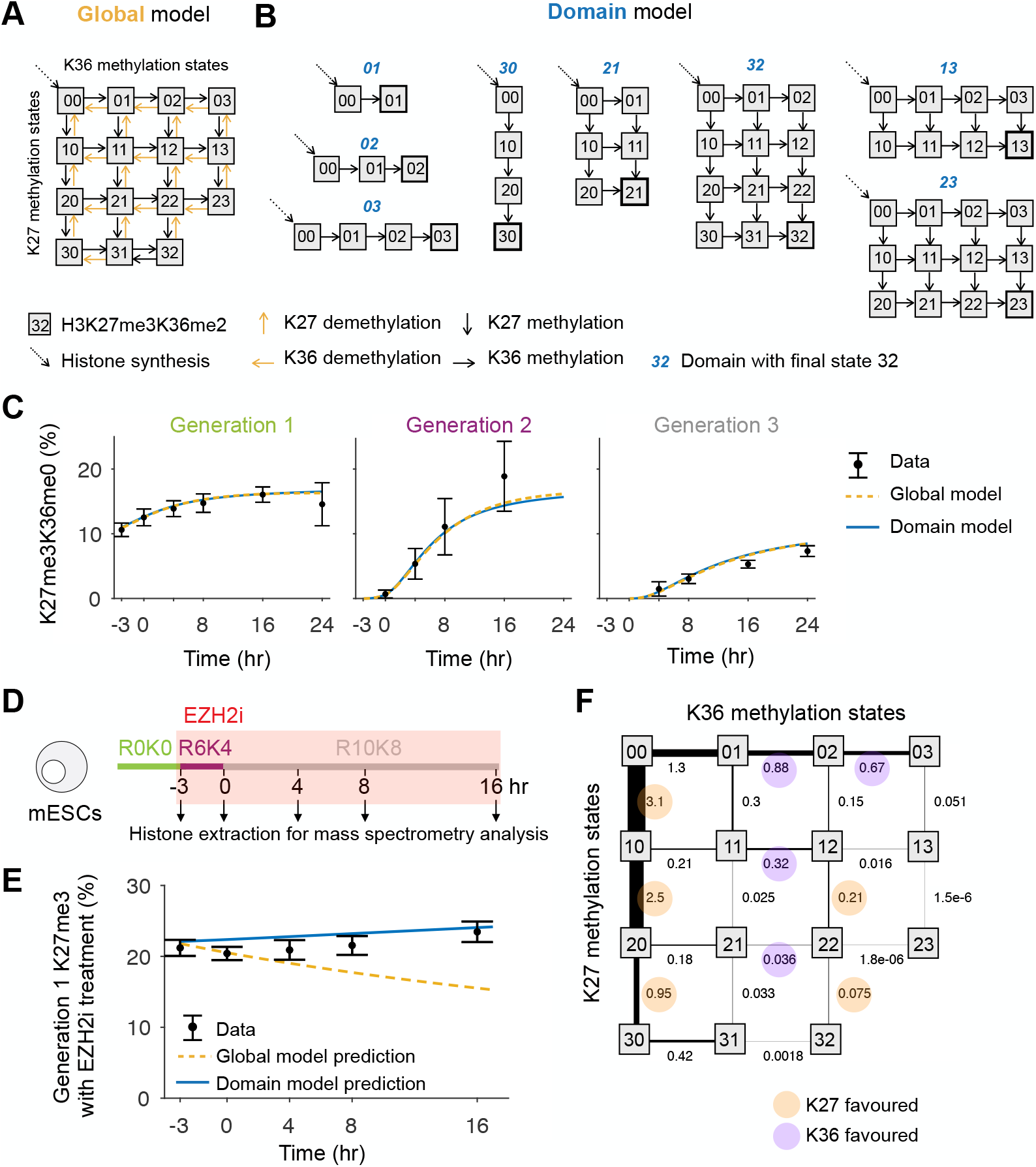
Computational domain model predicts dynamics during inhibitor treatment without global demethylation. **A.** The global model describes the abundance of methylations on K27 and K36 with methylation and demethylation. Differences between generations occur from the incorporation of unmodified histones (dotted diagonal arrow): This only occurs for the time where the cells are in the corresponding culture medium. **B.** The domain model assumes that the histone tails are methylated until they reach a defined final state, which depends on the domain the histone belongs to. In contrast to the global model, it requires no demethylation. **C.** Both the global and the domain model can describe methylation dynamics of untreated cells in all three generations. **D.** Experimental design of EZH2 inhibitor treatment. **E.** Only the domain model prediction fits to the K27me3 levels of generation 1 during inhibitor treatment. **F.** The model averaged fluxes for the population steady state show antagonism between K27 and K36 methylation. See also Figure S2.

To examine the existence of global demethylation as the testable difference of the two model hypotheses, we inhibit K27 methylation with an EZH2 inhibitor (EPZ-6438) and measure the histone modification dynamics over time (Fig. 2D). K27me3 levels remain stable at around 20% for 16 hours (Fig. 2E). We use this finding to test the two models. First, we estimate the efficiency of the EZH2 inhibitor from the ratio of K27me3 to K27me2 in untreated and treated cells to be around 92.8±5.2% (mean±std., n=3 replicates; see Fig. S2F, G and Supplementary Information for details). We then predict K27me3 levels upon inhibitor treatment for the two model hypotheses. The experimentally observed stability of K27me3 in the presence of the inhibitor is inconsistent with the global model that assumes demethylation, but can be well explained quantitatively with our domain model (Fig. 2E). We thus reject the global model and use the domain model to analyze our data further.

To interrogate interactions between the two residues, we calculate the methylation flux in steady state, defined as the product of the modification state abundance times the reaction rate constant (Supplementary Information). The fluxes from K27me0 to K27me3 with no K36 methylation make up 57.4% of all fluxes (Fig. 2F). We find evidence for antagonistic behavior in both directions: In all cases with fluxes over 0.5 hr^−1^, the addition of a methyl mark to the site with the higher modified state is more likely than modification of the less modified site (Fig. 2F and Fig. S2H). For example, there is a 2.5/0.21 = 11.9-fold increased probability for further methylation of K27 on a histone already carrying K27me1K36me0 as compared to methylation of K36, and a 0.67/0.15 = 4.5-fold increase for further methylation of K36 on K27me0K36me2 as compared to adding methylation to K27 (Supplementary Information). In cases where the lysines are equally methylated, K27 methylation is favored on naïve histones (K27me0K36me0) and on di-methylated histones (K27me2K36me2), while K36 methylation is favored when both residues are mono-methylated (K27me1K36me1). Collectively, our modelling provides a detailed, quantitative picture of the antagonistic relationship between K27me and K36me.

### Pre-existing K27me3 on old histones increase the rate of de novo K27me3

Based on the ability of K27me3 to allosterically activate PRC2, it has been predicted that parental histones carrying K27me3 directly stimulate K27me3 establishment on newly incorporated histones (Margueron et al., 2009; Sanulli et al., 2015). Our quantitative system yields the unique opportunity to test this hypothesis by directly measuring the contribution of parental K27me3 to K27me3 establishment on new histones (Fig. 3A). To erase K27me3 from parental histones, cells were treated 7 days with the EZH2 inhibitor (Fig. 3B) (Hojfeldt et al., 2018). After inhibitor treatment, the level of K27me3 dropped below 3% and K27me2 was also strongly reduced, while K27me1 accumulated (Fig. 3C). This is consistent with previous work (Hojfeldt et al., 2018) and suggests that in this setting EZH1, which is not targeted by the inhibitor, catalyzes K27me1. Focusing on the methylation kinetics on new histones (generation 2), we found that the establishment of K27me2/3 methylation was delayed in cells lacking K27me3 (Fig. 3D). Establishment of other methylation marks including H3K9me3 (Fig. S3A) was not affected, excluding unspecific effects on methylation dynamics. We performed quantitative model selection to identify rate constants that differ between generation 2 histones of untreated and recovering cells (Fig. 3E, S3B, Supplementary Information). We found that a model with substantial differences in K27 mono-methylation of K27me0K36me0, K27 tri-methylation of K27me2K36me0, and K36 mono-methylation of K27me0K36me0 and K27me2K36me0 is able to explain the data and the differences between untreated and recovering cells (Fig. 3E, S3B). As expected, the most important difference found by model selection is the K27 mono-methylation of K27me0K36me0, followed by the K27 tri-methylation of K27me2K36me0. Together, these results strongly support the existence of a feed-forward loop, whereby parental K27me3 stimulates establishment of K27 methylation on new histones by increasing the enzymatic rate.

**Figure 3.**
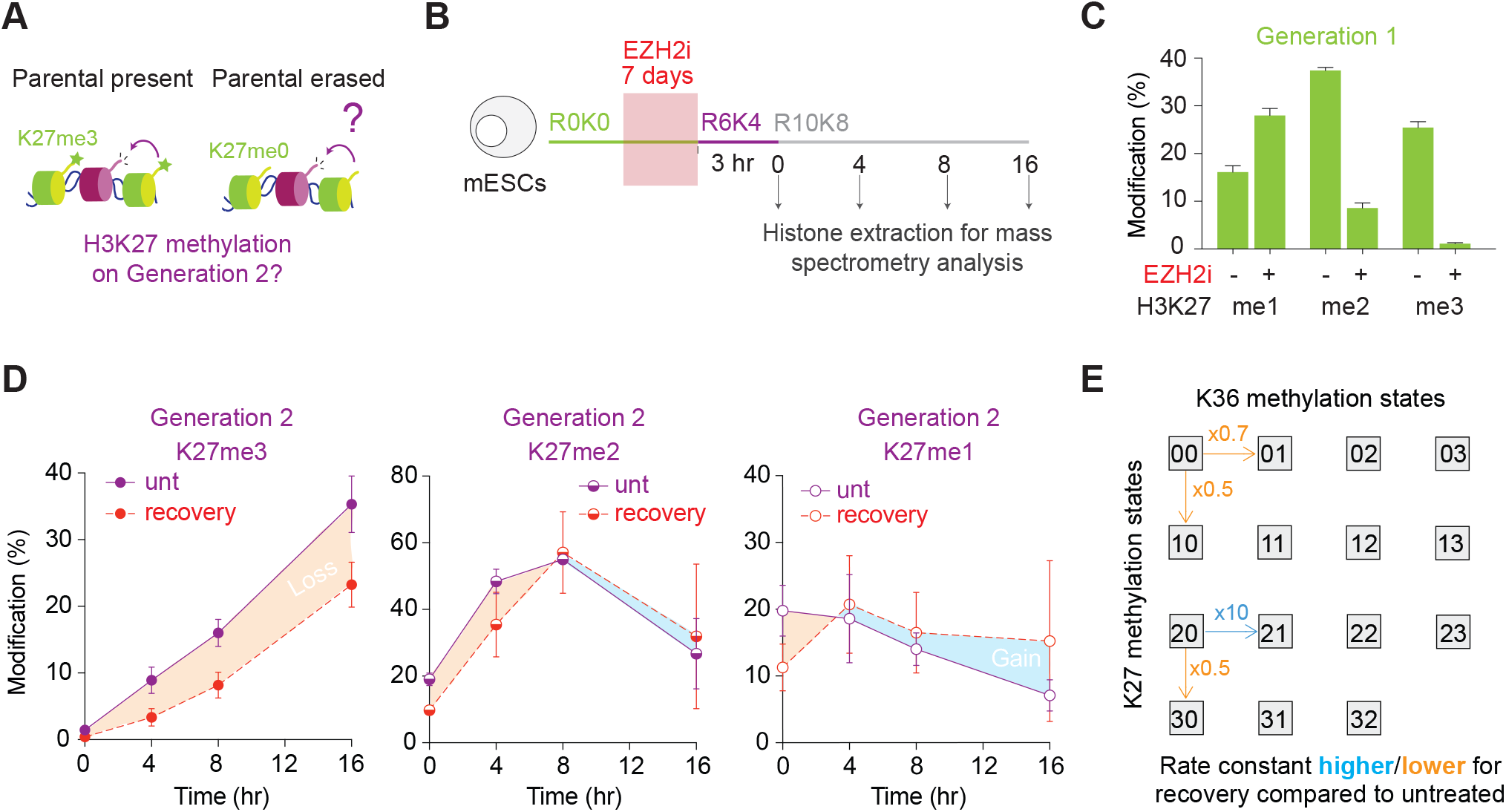
Lack of K27me3 on parental histones reduces the rate of de novo K27me3. **A.** Illustration of the question; do pre-existing K27me3 affect the kinetics of *de novo* K27me3 establishment? **B.** Experimental design to track *de novo* K27 methylation on generation 2 histones upon recovery from 7 days of EZH2 inhibitor treatment. **C.** K27 methylation levels on generation 1 histones after 7 days of EZH2 inhibitor treatment (measured at time 0 in B). **D.** K27 methylation dynamics on generation 2 histones with losses and gains in the absence of pre-existing K27me3 highlighted by orange and blue shaded areas, respectively. **E.** The domain model predicts differences in the rate constants between untreated and recovering generation 2 histones. See also Figure S3.

### K36 methylation can lock cells in an aberrant modification state

Cells fully restored K27me3 levels after removal of the inhibitor within approximately five generations (Fig. S3C), as reported previously (Hojfeldt et al., 2018). This relatively slow recovery was unlikely to only reflect the delay in K27me2/3 on new histones, manifested on a scale of hours rather than cell generations. We thus also analyzed the recovery of the old histones (generation 1) after inhibitor treatment, expecting them to rapidly gain K27me3. Surprisingly, a large proportion of generation 1 histones were refractory to K27me3 after inhibitor removal, as compared to new generation 2 histones present in the same cells (Fig. 4A). This result was counter intuitive given that a substantial fraction of generation 1 histones already carried K27me1 at the time of inhibitor removal (Fig. 3C), and we thus sought to identify the basis for this recovery defect.

**Figure 4.**
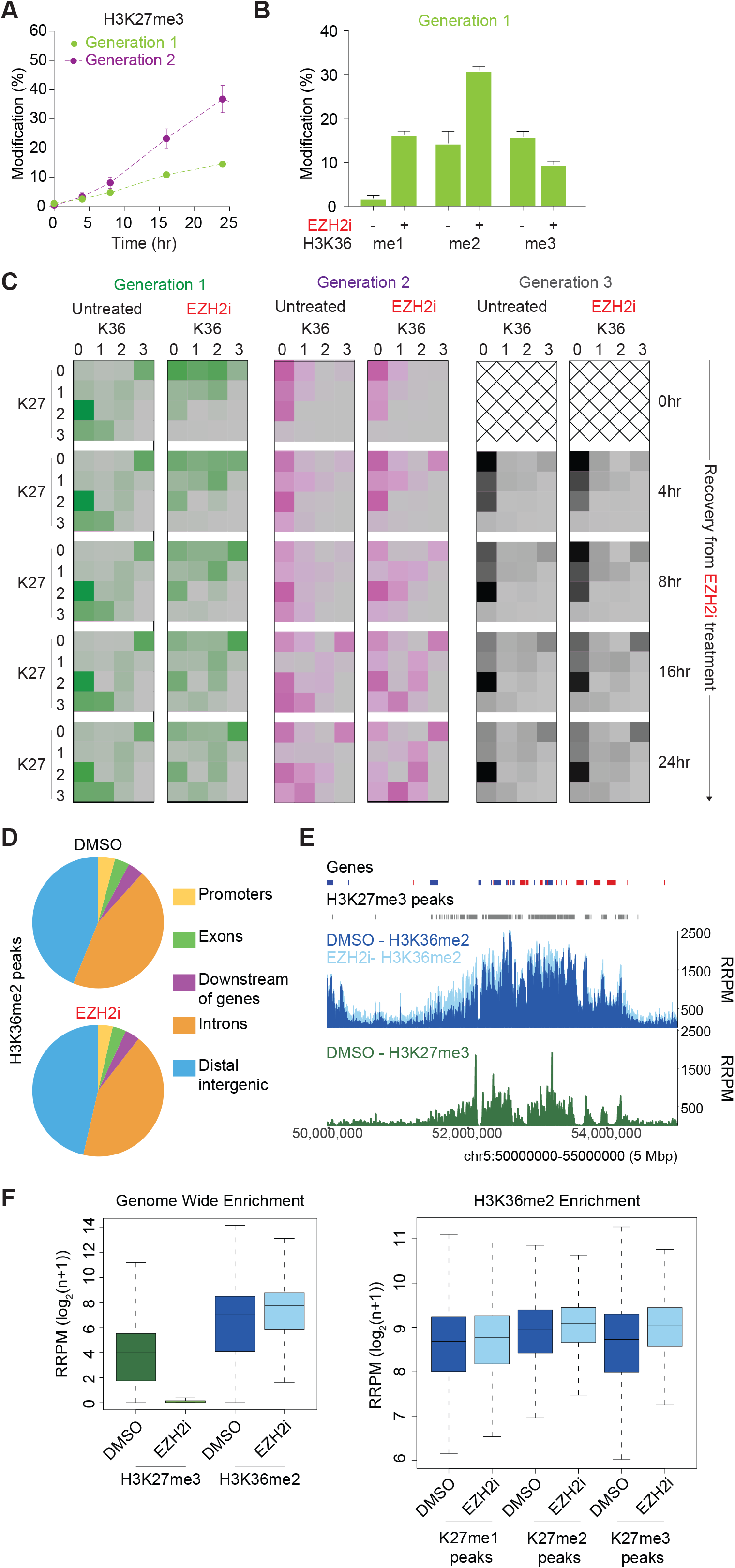
K36 methylation locks Polycomb domains in an aberrant modification state. **A.** K27me3 levels on generation 1 and generation 2 histones during recovery from EZH2 inhibitor treatment as described in Fig. 3B. **B.** K36 methylation levels on generation 1 histones after 7 days of EZH2 inhibitor treatment (time 0 in Fig. 3B). **C.** Heatmaps of K27 and K36 methylation levels on generation 1, 2 and 3 histones during recovery from inhibitor treatment described in Fig. 3B. **D.** Pie chart showing distribution of H3K36me2 peaks in different genomic features in DMSO or EZH2i treated samples. **E.** Screenshot of H3K36me2 ChIP-seq signal in DMSO (dark blue) or EZH2i (light blue) treated samples (top) and H3K27me3 ChIP-seq signal in DMSO treated samples (bottom, green). Quantitated with RRPM, log2(n + 1) over the region depicted. Red and blue blocks represent genes in the forward or reverse strand, respectively. Grey boxes indicate H3K27me3 peaks in untreated conditions. **F.** Left, boxplot of H3K36me2 and H3K27me3 ChIP-seq signal over 1kb windows across the genome in DMSO and EZH2i treated samples. Black line, median; dashed lines, 1.5 × interquartile range. Quantitated with RRPM, log2(n + 1). Right, boxplot of H3K36me2 ChIP-seq signal over 1kb windows across H3K27me1, H3K27me2 and H3K27me3 peaks. Black line, median; dashed lines, 1.5 × interquartile range. Quantitated with RRPM, log2(n + 1). See also Figure S4.

In depth analysis of generation 1 histones revealed that there was a strong skew towards more K36me1/2 after inhibitor treatment (Fig. 4B) as well as a moderate loss of K36me3. This is consistent with recent findings in EED knock-out mESCs, where loss of K27 methylation leads to a 3-fold increase in K36me2 and reduction of K36me3 (Oksuz et al., 2018). Similar effects on K36me were observed upon genetic inhibition of PRC2 by expression of H3K27M (Fig. S4A, B), excluding that K36 methylation changes was due to unspecific activity of the EZH2 inhibitor. Remarkable, our setup showed that the strong positive K36me1/2 skew was maintained throughout the lifetime of these histones, regardless that the EZH2 inhibitor was removed (Fig. 4C, generation 1).

K36me2/3 inhibits PRC2 mediated K27me2/3 in vitro (Schmitges et al., 2011; Yuan et al., 2011) and modelling analysis (Fig. 2F and (Zheng et al., 2012)) indicate that K36 methylation directly antagonizes K27 methylation. Taken together, this argues that K27me3 establishment on generation 1 histones is impaired because aberrant K36me2 inhibits PRC2 activity *in cis*. To further substantiate this hypothesis, we determined K36me2 occupancy across the genome by quantitative ChIP-seq analysis. Consistent with previous work (Streubel et al., 2018), K36me2 is a wide-spread mark in mESCs found in intergenic regions and across gene bodies (Fig. 4D, E) and also overlaps with K27me1/2/3 (Fig. S4F). This distribution of K36me2 was not dramatically changed upon long-term EZH2 inhibition (Fig. 4D, E). However, K36me2 levels increased genome-wide and in particular in regions normally enriched for K27me3 (Fig. 4F). Collectively, these results explain why K27me3 is impaired on generation 1 histones and reveals a surprising tardiness in reshaping the methylation landscape once established. Pre-existing K36 methylation makes cells refractory to change, allowing full recovery of K27me3 levels only after dilution of these pre-existing histones over 5 rounds of cell division. This underscores the importance of establishing correct histone modification patterns on new histones, as an unwarranted K36/K27 skew can affect cells across several cell divisions. Indeed, histones incorporated after removal of the inhibitor (generation 2 and 3) also showed reminiscence of the inhibitor induced K36 methylation skew (Fig. 4C, S4C-E). Aberrant K36me2 on parental histones thus impacted subsequent generations of histones that did not experience the inhibitor treatment, further underscoring that acquisition of modifications on new histones is influenced by the environment where they land.

## DISCUSSION

Understanding how histone-based information is propagated across cell division remains a major challenge in biology. Here we take a quantitative modelling-based approach to tackle this question. By measuring K27 and K37 methylation dynamics on distinct histones, we develop a computational model that is reusable and provides a new analysis framework to understand propagation of histone modifications. Our data-driven model selection rejects global K27 and K36 demethylation, provides in vivo rate constants for individual reactions and argues for the existence of distinct methylation domains. By studying recovery from EZH2 inhibition, we find that pre-existing K27me3 on old histones stimulates the rate of *de novo* K27me3 establishment. Our model also reveals a detailed, quantitative picture of the mutual antagonism between K27 and K37 methylation states. Consistent with this, inhibition of EZH2 reshapes the K36me landscape, which in turn stabilizes the aberrant state and impedes restoration of K27me3 upon inhibitor removal.

Our modelling approach is inspired by the model of Zheng et al. (2012). Indeed, the global model (Fig. 2A) has the same model structure as presented in Zheng et al (2012), although we have omitted 22 individual demethylation rates in favor of three demethylation rates for K27 and three for K36 to avoid overfitting. However, a model with demethylation cannot explain stable K27me3 states during inhibitor treatment: If K27me2 sites are no longer methylated, demethylation should lead to a clear decrease of K27me3 over time. We thus introduce a domain model (Fig. 2B) that is able to explain our data and that is consistent with no global demethylation (Reveron-Gomez et al., 2018). Our domain model argues for the existence of distinct populations of histones that progress towards specific states. This is consistent with well-established principles of chromatin organization where specific post-translational modifications occupy distinct genomic locations (CpG islands, gene bodies, promoter regions) and even form discrete 3D structures in the cell nucleus (Polycomb bodies). Thus, histones are not equal in their methylation potential, but their site of incorporation is decisive for the methylation state they will acquire. Similar to Zheng et al. (2012), we find antagonism between K27 and K36 methylation. While the authors there argue that this antagonism manifests in decreasing K36 methylation rate constants with increasing K27 methylations (and vice versa), we believe that a direct comparison of K27 and K36 methylation fluxes for a particular state is more appropriate. Indeed, once either K27 or K36 is set on an otherwise balanced histone, the addition of more methylation marks of the same kind is often more likely for high-flux states (Fig. 2F). Such a ‘the rich get richer’ analogy seems to be particularly relevant for understating the recovery from inhibition. There, histones that have traversed to a high K36 methylation state upon K27 inhibition are no longer able to regain untreated K27 levels, probably being stuck in a particular domain. Similar to other models that describe epigenetic modifications in a quantitative manner (Berry et al., 2017; Blasi et al., 2016), we also assume mass action kinetics for transitions between chromatin states. In contrast to other approaches, however, we explicitly fit combinatorial histone dynamics in a variety of experimental settings. This allows us to infer structure and kinetics in an unprecedented manner.

The intriguing read-write function of the enzymes that catalyze key repressive histone modifications like K27me3 and H3K9me3 argues that chromatin states are maintained across cell division through self-propagation (Reinberg and Vales, 2018). For PRC2 this read-write mechanism involves the recognition of K27me3 by an aromatic cage in the EED subunit, which in turn leads to allosteric activation of EZH2 (Reinberg and Vales, 2018). This mechanism is well understood, but its biological role remains a matter of debate. Self-propagation alone is not sufficient for K27me3 maintenance in Drosophila, as a genetic contribution from PREs is also required (Coleman and Struhl, 2017; Laprell et al., 2017). Establishment of the K27me3 landscape in mESCs can also occur in the absence of pre-existing K27me3 (Hojfeldt et al., 2018; Oksuz et al., 2018) and in EED allosteric activation mutants (Oksuz et al., 2018). However, EED allosteric activation mutants are delayed in K27 methylation and fails to re-establish wild-type levels of K27me3 (Oksuz et al., 2018), arguing for the importance of the read-write mechanism in K27me3 maintenance. However, PRC2 also methylates its binding partner JARID2 on a K27 mimicking peptide that can activate the enzyme allosterically (Sanulli et al., 2015) and this might contribute to reduced activity of the EED mutants. By measuring methylation on new histones specifically, we show that the lack of pre-existing K27me3 reduces the efficiency of de novo K27me3 establishment. Using mathematical modelling, we can assign this to a reduced rate of K27 methylation. This demonstrates that pre-existing H3K27me3 stimulates de novo K27 methylation during DNA replication and argues that self-propagation through a read-write mechanism is important for timely restoration of K27me3 domains.

Our model shows a complex pattern of mutual antagonism between methylation on K27 and K36. This corroborates a number of other *in vivo* and *in vitro* studies. *In vitro*, the presence of K36me2/3 reduces the activity of EZH2 mediated methylation in cis while there are no reports that K27me2/3 directly impair the activity of K36me2/3 methyltransferases (Jani et al., 2019; Schmitges et al., 2011; Yuan et al., 2011). In cancer cells expressing the H3K36M oncohistone, or upon NSD1 deletion in ESCs, inhibition of K36 methylation leads to aberrant gain and spreading of K27me3 (Lu et al., 2016; Streubel et al., 2018). Likewise, expression of the K27M oncohistone leads to increased levels of K36me2 and its aberrant spreading into K27me3 domains ((Stafford et al., 2018), this paper). Our modelling shows a two-way antagonism that is more complex and broadly involves K27 and K36 methylation stages. However, there is currently no evidence to support a direct effect of K27 methylation on K36 methyltransferases, leading us to suggest that this antagonism reflects chromatin compaction or reduced affinity of the K36 methyltransferases in a K27me context. Quantitative ChIP-seq analysis confirmed increased K36me2 upon EZH2 inhibition, but the effect was less dramatic than indicated by mass spectrometry analysis. Given that most nucleosomes show asymmetric K27me3 (Voigt et al., 2012), part of the K36me2 gain might thus reflect an increase in symmetric K36me that would be masked in ChIP-seq experiments. In cells, K27me3 and K36me3 domains do not generally overlap, but there is a substantial overlap of K36me2 signal with K27me2 occupancy as well as some overlap with K27me3 domains (this work, (Streubel et al., 2018)). It was suggested that these modifications might occur on different nucleosomes. However, mass spectrometry analysis shows substantial co-occurrence of these marks on the same histone tail (K36me2-3K27me1/2/3: 11,6%) (this work, (Jung et al., 2010; Zheng et al., 2012)), arguing that they are not mutually exclusive. Our modelling and kinetics analysis indicate that K27me is imposed first, followed by methylation of K36. This explains how histone H3 can acquire both marks despite K36me directly impairing PRC2 activity. If K36me is allowed to establish first (as in the context of EZH2 inhibition), this in turn impairs K27me2/3 establishment on those nucleosomes.

We propose that one biological implication of the K27/K36 methylation antagonism is to enhance stability of epigenetic states. Methylation on these sites, in particular K36me2 and K27me3 ((Streubel et al., 2018) and this work), are keeping each other in check by restraining unwarranted spreading. We observed that upon recovery from EZH2 inhibition, pre-existing histones did not acquire K27me3 and retained aberrant high level of K36me2. The epigenetic-state of old histones are thus locked, in this case in an aberrant state, and changing this state during recovery relies on dilution by new naive histones during replication in order to re-set the K27/K36 methylation balance. This resistance to change is further enhanced by the lack of EZH2 allosteric activation in early recovery (see above), and, intriguingly, a slight skew towards K36me for several histone generations. The prediction from these observations and the fact that almost all histone tails carry some level of K27/K36 methylation, is that the K27/K36 methylation antagonism stabilize epigenetic states and provide a barrier to changes in cell identity that can be overridden globally by cell proliferation. This is relevant for cancer etiology where the K27/K36 methylation balance is disturbed by mutations in both histones and chromatin regulators and targeted therapeutic intervention may reset the landscape.

## Supporting information

Supplemental Modeling

## Acknowledgments

We thank Jonas Hojfeldt and the K. Helin laboratory for advice on mESCs and EZH2 inhibitor treatment, and Kathleen Stewart-Morgan and members of the Groth, Imhof and Marr laboratories for fruitful discussion. This work was supported by a collaborative grant from the Lundbeck Foundation (R198-2015-269) to A.G., A.I and C.M. Research in the Groth lab was additionally supported by the European Research Council (ERC CoG no. 724436) and Independent Research Fund Denmark (7016-00042B). C.A. is supported by the European Research Council (ERC Stg no. 715127).

## Author Contributions

C.A. perform the cell work. M.VA performed all mass spectrometric measurements and analyzed the data together with C.A., C.L. and A.I. I.F. assisted with mass spectrometry. C.L., J.H. and C.M. developed the models. C.L. developed and L.S. reviewed the code. C.L. wrote the modelling supplement with L.S., C.M. and J.H. N.R-G. and S.G. performed ChIP-seq analysis. C.A. and A.G. wrote the manuscript with input from all authors.

## SUPPLEMENTARY INFORMATION

### Experimental Model and Subject Details

#### E14 mESCs cell culture

E14 mESCs were cultured at 37°C and 5% Co2 on gelatin-coated plates in 2i/LIF medium (2i custom made medium (Thermo Fisher)) supplemented with Pen-Strep (Gibco), 2mM Glutamax (Gibco), 50μM β-mercaptoethanol (Gibco), 0.1mM nonessential amino acids (Gibco), 1mM sodium pyruvate (Gibco), N2+B27 (Thermo Fisher), GSK3i (CHIR99021), MEKi (PD0325901), Leukemia Inhibitor Factor (LIF; produced in Kristian Helin laboratory). Medium was supplemented with light, medium or heavy arginine and lysine (R0 Arginine hydrochloride (Sigma A6969), R6 Arginine 13C6 (CNLM-2265-H1), R10 Arginine 13C6 15N4 (CNLM-539-H1), K0 Lysine hydrochloride (Sigma L8662), K4 Lysine 4,4,5,5-d4 (DLM-2640-1 1gram), K8 Lysine 13C6 15N2 (CNLM-291-H-1 1gram) Cambridge isotope). We passaged cells every 2-3 days by removing the medium, washing cells in PBS, dissociating cells with 0.25% trypsin EDTA (Gibco) with gentle disruption of colonies by pipetting, resuspending cells in medium, pelleting by centrifugation and resuspending cells and plating at density of 5×10^6^ cells/15cm dish.

#### Drosophila S2 Cell Culture

S2 cells were grown in suspension in spinners in M3+BPYE media: Shields and Sang M3 Insect Medium (Sigma, S-8398), KHCO3 (Sigma, 12602), yeast extract (Sigma, Y-1000), bactopeptone (BD, 211705), 10% heat-inactivated FCS (GE Hyclone, SV30160.03) and 1X penicillin/streptomycin (GIBCO, 151400122). Cells were incubated at 25°C with 5% CO2.

### Method Details

#### EZH2 inhibition and wash out

E14 mESCs were treated with 10μM of EPZ6438 EZH2 inhibitor (MedChem Express) for indicated time. For washout, cells were washed three times in PBS, trypsinized, washed in growth medium, and plated into R6K4 fresh medium.

#### Sample preparation for histone modification analysis by MS

Acid extracted histones were resuspended in Lämmli buffer and separated by a 14-20% gradient SDS-PAGE, stained with Coomassie (Brilliant blue G-250). Protein bands in the molecular weight range of histones (15-23 kDa) were excised as single band/fraction. Gel slices were destained in 50% acetonitrile/50mM ammonium bicarbonate. Lysine residues were chemically modified by propionylation for 30min at RT with 2.5% propionic anhydride (Sigma) in ammonium bicarbonate, pH 7.5 to prevent tryptic cleavage. This step added a propionyl group only to unmodified and monomethylated lysines, whereas lysines with other side chain modification will not obtain an additional propionyl-group. Subsequently, proteins were digested with 200ng of trypsin (Promega) in 50mM ammonium bicarbonate overnight and the supernatant was desalted by C18-Stagetips (reversed-phase resin) and carbon Top-Tips (Glygen) according to the manufacturer’s instructions. Following carbon stage tip, the dried peptides were resuspended in 17μl of 0.1% TFA.

#### LC-MS analysis of histone modifications

5μl of each sample were separated on a C18 home-made column (C18RP Reposil-Pur AQ, 120 × 0.075mm × 2.4μm, 100Å, Dr. Maisch, Germany) with a gradient from 5% B to 30% B (solvent A 0.1% FA in water, solvent B 80% ACN, 0.1% FA in water) over 32min at a flow rate of 300nl/min (Ultimate 3000 RSLC Thermo-Fisher, San Jose, CA) and directly sprayed into a Q-Exactive HF mass spectrometer (Thermo-Fisher Scientific). The mass spectrometer was operated in the PRM mode to identify and quantify specific fragment ions of N-terminal peptides of human histone 3.1 and histone 4 proteins. In this mode, the mass spectrometer automatically switched between one survey scan and 9 MS/MS acquisitions of the m/z values described in the inclusion list containing the precursor ions, modifications and fragmentation conditions (Supplemental Table 1). Survey full scan MS spectra (from m/z 250–800) were acquired with resolution 30,000 at m/z 400 (AGC target of 3×106). PRM spectra were acquired with resolution 15,000 to a target value of 2×105, maximum IT 60ms, isolation 2 window 0.7m/z and fragmented at 27% normalized collision energy. Typical mass spectrometric conditions were: spray voltage, 1.5kV; no sheath and auxiliary gas flow; heated capillary temperature, 250°C.

#### Quantification of histone modifications

Data analysis was performed with Skyline (version 3.6) (MacLean et al., 2010) by using doubly and triply charged peptide masses for extracted ion chromatograms (XICs). Peaks were selected manually and the integrated peak values (Total Area MS1) were exported as .csv file for further calculations. The percentage of each modification within the same peptide is derived from the ratio of this structural modified peptide to the sum of all isotopically similar peptides. Therefore, the Total Area MS1 value was used to calculate the relative abundance of an observed modified peptide as percentage of the overall peptide. The unmodified peptide of histone 3.1 (aa 41–49) was used as indicator for total histone 3.1. Coeluting isobaric modifications were quantified using three unique MS2 fragment ions. Averaged integrals of these ions were used to calculate their respective contribution to the isobaric MS1 peak (e.g., H3K36me3 and H3K27me2K36me1). The mass spectrometry proteomics data have been deposited to the ProteomeXchange Consortium via the PRIDE partner repository with the dataset identifier PXD014807.

#### ChIP-seq

Cells were grown for 7 days in 2i/LIF media supplemented with 10μM EZH2i (EPZ6438) or equivalent volume of DMSO. Cells were then processed using the truChIP Chromatin Shearing Kit (Covaris, 520127). In brief, cells were washed and fixed in 1% formaldehyde for 5 minutes. Crosslinking was quenched by adding Covaris quenching buffer and the reaction was incubated for 5 minutes. Fixed cells were washed twice in 1X PBS, scraped off in 1X PBS and spun down. Cell pellets were snap-frozen in liquid nitrogen, and stored at −80°C until lysis. Nuclei isolation was performed on fixed cells following manufacturer’s instructions. 20 million cells were sonicated in 1mL tubes using a Covaris M220 with the following settings: duty cycle 10% intensity, 200 cycles/ burst, 20 minutes processing time, 7°C bath temperature, water level full. Sonicated chromatin was centrifuged at 14,000 rpm at 4°C for 10 minutes and the supernatant was isolated for subsequent steps. In parallel, Drosophila S2 cells were fixed, lysed and sonicated as described above. For quantitative ChIP analysis of H3K36me2 and H3K27me3 input chromatin was mixed with Drosophila S2 chromatin (0.05% of total chromatin) after sonication. 30μg of mESCs sonicated chromatin or 30μg total of mixed mESCs and Drosophila S2 sonicated chromatin were diluted up to 500mL with dialysis buffer (4% glycerol, 10mM Tris-HCl, 1mM EDTA, 0.5mM EGTA; pH 8) and 400mL of incubation buffer (2.5% Triton X-100, 0.25% sodium deoxycholate, 0.25% SDS, 0.35M NaCl, 10mM Tris-HCl; pH 8) supplemented with leupeptin, aprotinin, pepstatin, and PMSF. Chromatin was pre-cleared with either Protein G agarose beads pre-coupled with bridging antibody (Active Motif #53017) following manufacturer’s instruction (pG-Ab) or Protein A agarose beads for 1 hour at 4°C. After pre-clearing, chromatin was incubated overnight at 4°C with 10μg of the appropriate antibody (H3K27me1: Active Motif mAb #61015; H3K27me2: Cell Signalling mAb #9728; H3K27me3: Cell Signaling mAb #9733; H3K36me2: AbCam mAb #176921), followed by incubation for 3 hours with either pre-blocked Protein G-Ab or Protein A agarose beads (incubated in 1 mg/ml BSA in RIPA buffer overnight). Chromatin bound to beads was washed three times in RIPA buffer (140 mM NaCl, 10mM Tris-HCl, 1mM EDTA, 1% Triton X-100, 0.1% SDS, 0.1% sodium deoxycholate, 1mM PMSF; pH 8), four times in RIPA buffer with 0.5 M NaCl, once in LiCl buffer 3 (250mM LiCl, 10mM Tris-HCl, 1mM EDTA, 0.5% NP-40, 0.5% sodium deoxycholate; pH 8) and twice in TE (10mM Tris-HCl, 1mM EDTA; pH 8). Chromatin was incubated with 125μg/ml RNase A for 30 minutes at 37°C SDS was then added to a final concentration of 1% and samples were incubated with 250μg/ml proteinase K for 10 hours at 37°C followed by 6 hours incubation at 65°C for de-crosslinking. De-crosslinked DNA was purified and size selected with Agencourt AMPure XP beads (Beckman Coulter, A63881) by using first a 0.55:1 ratio followed by a 3:1 final ratio to obtain fragments between 200-700 bp. Finally, 10ng of purified DNA was subjected to end repair, A-tailing and amplification using the KAPA Hyperprep kit protocol (Roche, KK8504). Before and after amplification (7 PCR cycles) DNA was cleaned-up with Agencourt AMPure XP beads at a 0.8:1 ratio.

#### Data sequencing and processing

ChIP-seq libraries were sequenced 75 bp single-end on an Illumina NextSeq 500. Trim Galore was used to trim adapter sequences. Reads were mapped to the mm10 assembly mouse genome using Bowtie2 (Langmead and Salzberg, 2012). Reads with MAPQ < 20, PCR duplicates, and reads that overlapped with the Broad Institute sequencing blacklist (Consortium, 2012) were filtered out. For downstream analyses, remaining reads were extended by 250bp. Reads were mapped to the dm3 assembly Drosophila genome using Bowtie2 (Langmead and Salzberg, 2012) to calculate reference-adjusted reads per million (RRPM) normalization factors. The number of uniquely mapped reads after deduplication was used to calculate RRPM as in (Reveron-Gomez et al., 2018).

#### Peak calling

Peak calls were created with SICER (Zang et al., 2009) using redundancy threshold = 1, window size = 500, fragment size = 250, effective genome fraction = 0.79, gap size = 1500 and an FDR threshold of 0.01. Input samples were used in peak calling to assess background and to determine true signal from IP enrichment.

#### Quantification and statistical analysis

Bedgraphs for screenshots were generated and visualized using Seqmonk (version 1.42.1). Boxplots were generated in R using custom scripts. Pie charts for annotation of genomic features were generated using CEAS (Shin et al., 2009).

#### Mathematical modeling of histone tail methylation

Due to format restriction, this section is in a separated document, Supplemental_Modeling.pdf

### Quantification and Statistical Analysis

Figure 1, 6<n<9 biological replicates have been performed. Figure 2D, 3 biological replicates have been performed. Figure 3 and 4A-C, 3 biological replicates have been performed. Figure 4D-F, 2 biological replicates have been performed. The mean with standard deviation is shown.

### Data and Code Availability

The mass spectrometry dataset generated during this study is available at Pride [PXD014807]. The sequencing dataset generated during this study is available at GEO [GSE135029]. The MATLAB code used for the mechanistic modelling is available at Zenodo [https://doi.org/10.5281/zenodo.3353481].

