## Supplemental Figures for "Domain model explains propagation dynamics and stability of K27 and K36 methylation landscapes"

**SUPPLEMENTARY FIGURE LEGENDS**

**Figure S1, related to Figure 1. A.** Proportion of each histone generations across the time course, based on quantification of the H3 aa41-49 peptide that remains unmodified. The proportion is given as a percentage such that the sum equals 100 % at any given time. **B.** Total modification levels across the time course, represented by pre-existing histones (green), combinations of the different generations (blue) and third generation histones (grey). For each modification, the sum is given (e.g. K27me3 is given as the sum of K27me3K36me0, K27me3K36me1, K27me3K36me2, and K27me3K36me3). **C, D.** Methylation levels on generation 2 histones for K27 and K36, on all individual peptides used to generate the sum shown in Fig. 1D. **E.** Methylation levels on histone generations 1, 2 and 3 for H3K9. (un) unmodified; (me1) mono-methylated; (me2) di-methylated; (me3) tri-methylated. **F.** Heatmap showing stepwise methylation of H4K20 methylation on generation 2 histones. (un) Unmodified; (me1) mono-methylated; (me2) di-methylated; (me3) tri-methylated.

**Figure S2, related to Figure 2. A.** Comparing different offsets by which data and model simulation are shifted shows highest QQ correlations for 10-1. **B.** Correlation for the QQ-plots for varying offsets. The maximum correlation is achieved for offset $10^{-1}$. **C.** Model fits including the global model without K27 demethylation for K27me3K36me0 and K27me3K36me1. The models were calibrated for the full data set. **D.** Both the global and the domain model is able to fit methylation dynamics in all generations. **E.** Model reduction for the domain model. The difference in BIC values is shown for the best tested model with given numbers of domains. A model with 8 domains has the best Bayesian Information Criterion (BIC) value, and there are models with 7 and 9 domains which cannot be rejected according to their BIC value. The dotted line shows the threshold of $\Delta$BIC=10. **F.** Illustration of the parametrization for the efficiency $\kappa$ of the inhibitor. **G.** Estimates for $\kappa$ comparing K27me3 and K27me2 levels for generation 3 histones for untreated cells and cells with EZH2 inhibitor. **H.** Fluxes obtained with the maximum likelihood estimates (MLEs) for all calibrated models. The models are color coded according to their BIC value. The model averaged (using BIC weights) fluxes are highlighted with red and fluxes which are compared in Fig. 2F are highlighted by arrows.

**Figure S3, related to Figure 3. A.** H3K9me3 levels on generation 2 and 3 histones. Experimental design as described in Fig. 3B. **B.** K27 and K36 methylation levels on generation 2 histones for untreated cells and during recovery. Experimental measurements and the model fit allowing for differences in the methylation rate constants $k_{00\text{->}01}$, $k_{00\text{->}10}$, $k_{20\text{->}21}$ and $k_{20\text{->}30}$ are shown. **C.** Total methylation levels on K27 across the time course on the combination of all three histone generations calculated as in Fig. S1B. Losses and gains compared to untreated control are by pink and blue shaded areas, respectively.

**Figure S4, related to Figure 4. A.** Bar-diagram showing K27 and K36 methylation levels in TIG-3 cells expressing H3K27M and H3K9M and untransduced controls (Ctrl). **B.** Heatmap showing K27 and K36 methylation levels in TIG-3 cells expressing H3K27M compare to control (Ctrl). **C-E.** K36 methylation levels on generation 2 histones in untreated conditions and upon recovery from EZH2 inhibitor treatment. **F.** Scatterplots showing direct comparison of ChIP-seq signal over 1kb windows across the genome for the indicated pairs of histone modifications. Quantitated with RPM, log2(n+1). Pearson's correlation R value for each plot is shown.

#### Figure S1

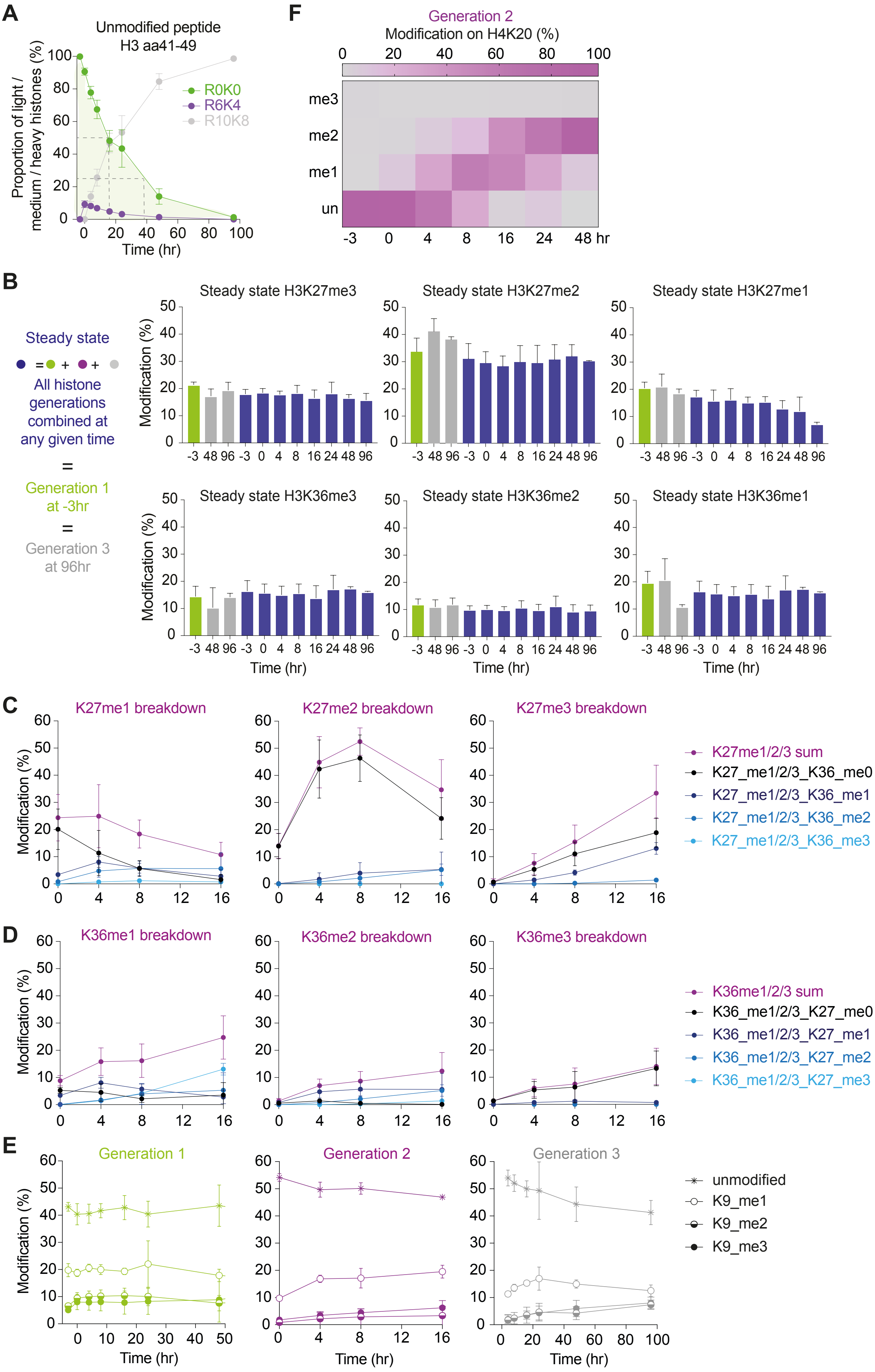
A
Unmodified peptide
H3 aa41-49
Proportion of light / medium / heavy histones (%)
R0K0
R6K4
R10K8
Time (hr)
F
Generation 2
Modification on H4K20 (%)
me3
me2
me1
un
-3 0 4 8 16 24 48 hr
B
Steady state
All histone generations combined at any given time
=
Generation 1 at -3hr
=
Generation 3 at 96hr
Steady state H3K27me3
Steady state H3K27me2
Steady state H3K27me1
Steady state H3K36me3
Steady state H3K36me2
Steady state H3K36me1
Modification (%)
Time (hr)
C
K27me1 breakdown
K27me2 breakdown
K27me3 breakdown
K27me1/2/3 sum
K27_me1/2/3_K36_me0
K27_me1/2/3_K36_me1
K27_me1/2/3_K36_me2
K27_me1/2/3_K36_me3
D
K36me1 breakdown
K36me2 breakdown
K36me3 breakdown
K36me1/2/3 sum
K36_me1/2/3_K27_me0
K36_me1/2/3_K27_me1
K36_me1/2/3_K27_me2
K36_me1/2/3_K27_me3
E
Generation 1
Generation 2
Generation 3
unmodified
K9_me1
K9_me2
K9_me3
Modification (%)
Time (hr)

#### Figure S2

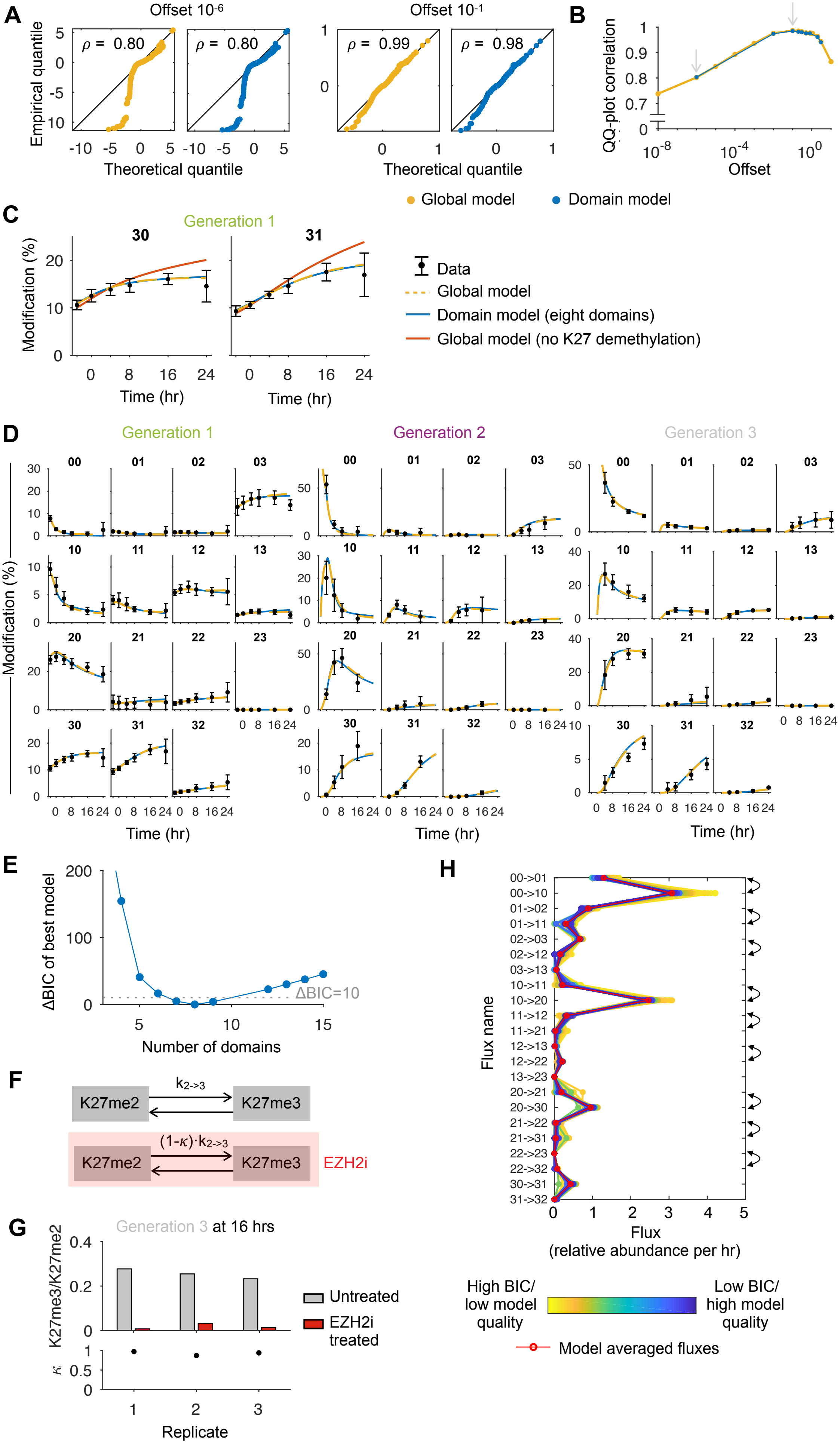
A
Offset 10-6
Offset 10-1
Empirical quantile
ρ = 0.80
ρ = 0.80
ρ = 0.99
ρ = 0.98
Theoretical quantile
Theoretical quantile
B
QQ-plot correlation
Offset
Global model
Domain model
C
Generation 1
30
31
Modification (%)
Time (hr)
Time (hr)
Data
Global model
Domain model (eight domains)
Global model (no K27 demethylation)
D
Generation 1
Generation 2
Generation 3
00
01
02
03
10
11
12
13
20
21
22
23
30
31
32
Modification (%)
Time (hr)
Time (hr)
Time (hr)
E
ΔBIC of best model
ΔBIC=10
Number of domains
F
K27me2
$k_{2->3}$
K27me3
K27me2
(1-κ)·$k_{2->3}$
K27me3
EZH2i
G
Generation 3 at 16 hrs
K27me3/K27me2
Untreated
EZH2i treated
κ
Replicate
H
Flux name
00->01
00->10
01->02
01->11
02->03
02->12
03->13
10->11
10->20
11->12
11->21
12->13
12->22
13->23
20->21
20->30
21->22
21->31
22->23
22->32
30->31
31->32
Flux
(relative abundance per hr)
High BIC/ low model quality
Low BIC/ high model quality
Model averaged fluxes

#### Figure S3

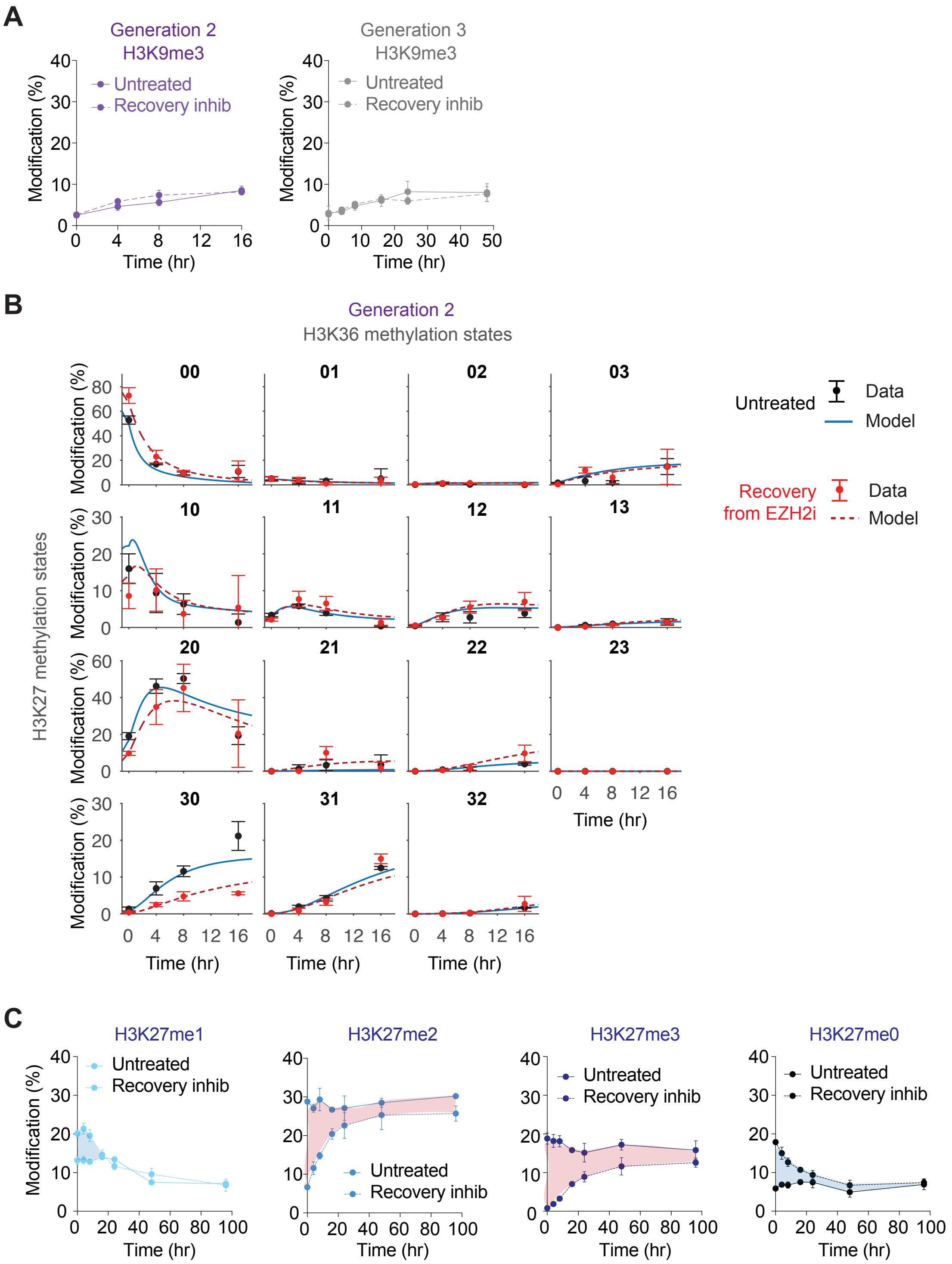

A
Generation 2
H3K9me3
Modification (%)
Untreated
Recovery inhib
Time (hr)
Generation 3
H3K9me3
Untreated
Recovery inhib
B
Generation 2
H3K36 methylation states
H3K27 methylation states
00
01
02
03
10
11
12
13
20
21
22
23
30
31
32
Untreated
Data
Model
Recovery
from EZH2i
Data
Model
C
H3K27me1
H3K27me2
H3K27me3
H3K27me0
Untreated
Recovery inhib

### Figure S4

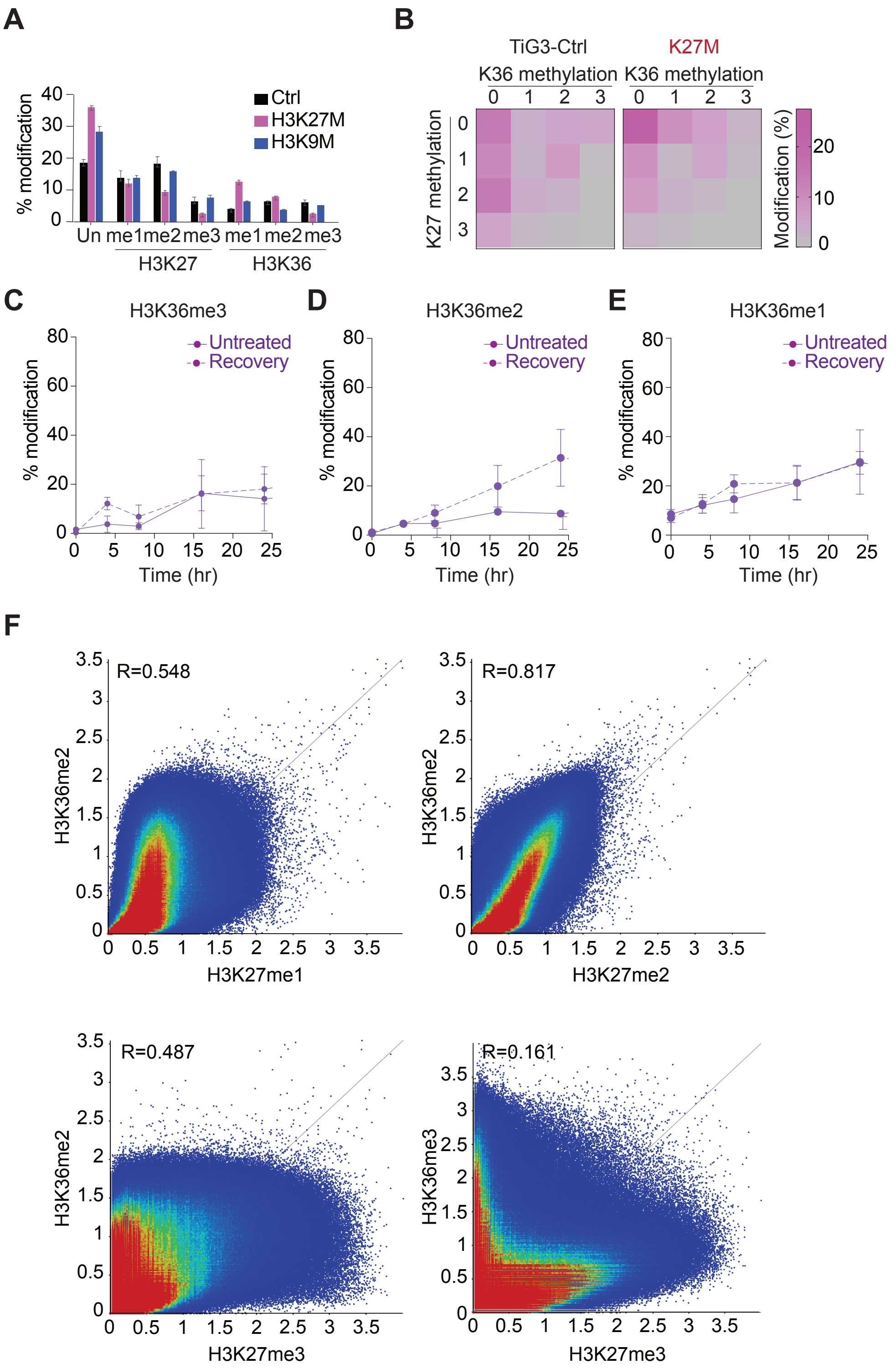
A
% modification
Ctrl
H3K27M
H3K9M
Un me1 me2 me3 me1 me2 me3
H3K27
H3K36
B
TiG3-Ctrl
K27M
K36 methylation
K27 methylation
Modification (%)
C
H3K36me3
Untreated
Recovery
Time (hr)
D
H3K36me2
E
H3K36me1
F
R=0.548
R=0.817
R=0.487
R=0.161
H3K36me2
H3K36me3
H3K27me1
H3K27me2
H3K27me3
