## Supplemental Modeling for "Domain model explains propagation dynamics and stability of K27 and K36 methylation landscapes"

### SUPPLEMENTARY INFORMATION
### Mathematical modeling of histone tail methylation

The dynamics of combinatorial histone modifications of H3K27 and H3K36 have been modeled previously with systems of ordinary differential equations based on the assumption of mass action kinetics (Zheng et al., 2012). Here, we also employ ordinary differential equation models based on mass action kinetics and describe the observed modifications, which are averaged over all cells and histones of the individual generations (Fig. 1A). We first describe the global model which includes demethylation, and then propose a different model that accounts for domains of specific modifications and that is able explain data obtain in an inhibitor experiment (Fig. 2D).

#### 1 Global model

The first model we considered consists of 45 state variables, with 15 state variables for each of the three generations (all possible combinations except K27me3K36me3) (Fig. 2A). It describes the change in modifications due to methylation and demethylation as well as dilution, which occurs when the cells divide and new, unmodified histones are incorporated. This model is similar to the one proposed by Zheng et al. (2012). To obtain a model for the relative abundance of modifications, we first derived the model for the absolute number of histone modifications.

The ODE system for the absolute number of histone modifications $\tilde{\mathbf{x}}_g = (\tilde{x}_{g,00}, \ldots, \tilde{x}_{g,23})$ reads for generation $g = 1, 2, 3$

$$\begin{aligned} \dot{\tilde{x}}_{g,ij} = {} & \chi_{\{(i,j)=(0,0)\}}(i,j)\, c_g(t) N \\ & + \chi_{\{i>0 \wedge (i,j)\neq(3,3)\}}(i,j)\, k_{i-1\,j \to i\,j}\, \tilde{x}_{g,i-1\,j} \\ & + \chi_{\{j>0 \wedge (i,j)\neq(3,3)\}}(i,j)\, k_{ij-1 \to ij}\, \tilde{x}_{g,ij-1} \\ & - \chi_{\{i<3 \wedge (i,j)\neq(2,3)\}}(i,j)\, k_{ij \to i+1\,j}\, \tilde{x}_{g,ij} \\ & - \chi_{\{j<3 \wedge (i,j)\neq(3,2)\}}(i,j)\, k_{ij \to ij+1}\, \tilde{x}_{g,ij} \\ & + \chi_{\{i<3 \wedge (i,j)\neq(2,3)\}}(i,j)\, d_{K27,i+1}\, \tilde{x}_{g,i+1\,j} \\ & + \chi_{\{j<3 \wedge (i,j)\neq(3,2)\}}(i,j)\, d_{K36,j+1}\, \tilde{x}_{g,ij+1} \\ & - \chi_{\{i>0 \wedge (i,j)\neq(3,3)\}}(i,j)\, d_{K27,i}\, \tilde{x}_{g,ij} \\ & - \chi_{\{j>0 \wedge (i,j)\neq(3,3)\}}(i,j)\, d_{K36,j}\, \tilde{x}_{g,ij}\,, \\ \dot{N} = {} & cN\,. \end{aligned} \tag{1}$$

The indicator function is denoted by $\chi$, $k_{ij\to i+1j}$ the rate constant for K27 methylation, $k_{ij\to ij+1}$ the rate constant for K36 methylation, $i$ denotes the number of methyl groups at K27, $j$ the number of methyl groups at K36, $N(t) = \exp(c(t - t_0))N_0$ the total number of histone tails with $N(t_0) = N_0$ the number of histone tails at the beginning of the experiment and $c$ being the cell division rate. Furthermore, $d_{K27,i}$ is the rate constant for demethylation, i.e., reducing the number of methyl groups at K27 from $i$ to $i - 1$. The special cases with $(i, j) \neq (2, 3)$ and $(i, j) \neq (3, 2)$ arise because we did not observe any K27me3K36me3 methylations. We model explicitely the three different generations of histones (Fig. 1A). Newly incorporated histones are unmodified and belong to the generation of the corresponding culture medium (Fig. 1A):

$$c_g(t) = \begin{cases} \chi_{\{t<-3\,\mathrm{h}\}}(t)\cdot c & g = 1\,, \\ \chi_{\{-3\,\mathrm{h}\leq t<0\,\mathrm{h}\}}(t)\cdot c & g = 2\,, \\ \chi_{\{t\geq 0\,\mathrm{h}\}}(t)\cdot c & g = 3\,, \end{cases} \tag{2}$$

The cell division rate $c$ is multiplied with the number of histone tails in (Eq. 1), because the number of histone tails is proportional to the number of cells and thus duplicated at cell division.

When changing the culture medium, initially no histones of this generation are present:

$$\tilde{x}_{g,ij}(t) = 0 \text{ for } \begin{cases} t < t_0 & g = 1\,, \\ t < -3\,\mathrm{h} & g = 2\,, \\ t < 0\,\mathrm{h} & g = 3\,, \end{cases} \qquad \forall i, j\,. \tag{3}$$

The full ODE system for (Eq. 1) reads

$$\begin{pmatrix} \dot{\tilde{x}}_{g,00} \\ \dot{\tilde{x}}_{g,01} \\ \dot{\tilde{x}}_{g,02} \\ \dot{\tilde{x}}_{g,03} \\ \dot{\tilde{x}}_{g,10} \\ \dot{\tilde{x}}_{g,11} \\ \dot{\tilde{x}}_{g,12} \\ \dot{\tilde{x}}_{g,13} \\ \dot{\tilde{x}}_{g,20} \\ \dot{\tilde{x}}_{g,21} \\ \dot{\tilde{x}}_{g,22} \\ \dot{\tilde{x}}_{g,23} \\ \dot{\tilde{x}}_{g,30} \\ \dot{\tilde{x}}_{g,31} \\ \dot{\tilde{x}}_{g,32} \end{pmatrix} = \begin{pmatrix}
1 & 0 & 0 & 0 & 0 & 0 & 0 & 0 & 0 & 0 & 0 & 0 & 0 & 0 & 0 \\
-1 & 1 & 0 & 0 & 0 & 0 & 0 & 0 & 0 & 0 & 0 & 0 & 0 & 0 & 0 \\
-1 & 0 & 0 & 0 & 1 & 0 & 0 & 0 & 0 & 0 & 0 & 0 & 0 & 0 & 0 \\
0 & -1 & 1 & 0 & 0 & 0 & 0 & 0 & 0 & 0 & 0 & 0 & 0 & 0 & 0 \\
0 & -1 & 0 & 0 & 0 & 1 & 0 & 0 & 0 & 0 & 0 & 0 & 0 & 0 & 0 \\
0 & 0 & -1 & 1 & 0 & 0 & 0 & 0 & 0 & 0 & 0 & 0 & 0 & 0 & 0 \\
0 & 0 & -1 & 0 & 0 & 0 & 1 & 0 & 0 & 0 & 0 & 0 & 0 & 0 & 0 \\
0 & 0 & 0 & -1 & 0 & 0 & 0 & 1 & 0 & 0 & 0 & 0 & 0 & 0 & 0 \\
0 & 0 & 0 & 0 & -1 & 1 & 0 & 0 & 0 & 0 & 0 & 0 & 0 & 0 & 0 \\
0 & 0 & 0 & 0 & -1 & 0 & 0 & 0 & 1 & 0 & 0 & 0 & 0 & 0 & 0 \\
0 & 0 & 0 & 0 & 0 & -1 & 1 & 0 & 0 & 0 & 0 & 0 & 0 & 0 & 0 \\
0 & 0 & 0 & 0 & 0 & -1 & 0 & 0 & 0 & 1 & 0 & 0 & 0 & 0 & 0 \\
0 & 0 & 0 & 0 & 0 & 0 & -1 & 1 & 0 & 0 & 0 & 0 & 0 & 0 & 0 \\
0 & 0 & 0 & 0 & 0 & 0 & -1 & 0 & 0 & 0 & 1 & 0 & 0 & 0 & 0 \\
0 & 0 & 0 & 0 & 0 & 0 & 0 & -1 & 0 & 0 & 0 & 1 & 0 & 0 & 0 \\
0 & 0 & 0 & 0 & 0 & 0 & 0 & 0 & -1 & 1 & 0 & 0 & 0 & 0 & 0 \\
0 & 0 & 0 & 0 & 0 & 0 & 0 & 0 & -1 & 0 & 0 & 0 & 1 & 0 & 0 \\
0 & 0 & 0 & 0 & 0 & 0 & 0 & 0 & 0 & -1 & 1 & 0 & 0 & 0 & 0 \\
0 & 0 & 0 & 0 & 0 & 0 & 0 & 0 & 0 & -1 & 0 & 0 & 0 & 1 & 0 \\
0 & 0 & 0 & 0 & 0 & 0 & 0 & 0 & 0 & 0 & -1 & 1 & 0 & 0 & 0 \\
0 & 0 & 0 & 0 & 0 & 0 & 0 & 0 & 0 & 0 & -1 & 0 & 0 & 0 & 1 \\
0 & 0 & 0 & 0 & 0 & 0 & 0 & 0 & 0 & 0 & 0 & 0 & -1 & 1 & 0 \\
0 & 0 & 0 & 0 & 0 & 0 & 0 & 0 & 0 & 0 & 0 & 0 & 0 & -1 & 1 \\
1 & 0 & 0 & 0 & -1 & 0 & 0 & 0 & 0 & 0 & 0 & 0 & 0 & 0 & 0 \\
0 & 1 & 0 & 0 & 0 & -1 & 0 & 0 & 0 & 0 & 0 & 0 & 0 & 0 & 0 \\
0 & 0 & 1 & 0 & 0 & 0 & -1 & 0 & 0 & 0 & 0 & 0 & 0 & 0 & 0 \\
0 & 0 & 0 & 1 & 0 & 0 & 0 & -1 & 0 & 0 & 0 & 0 & 0 & 0 & 0 \\
0 & 0 & 0 & 0 & 1 & 0 & 0 & 0 & -1 & 0 & 0 & 0 & 0 & 0 & 0 \\
0 & 0 & 0 & 0 & 0 & 1 & 0 & 0 & 0 & -1 & 0 & 0 & 0 & 0 & 0 \\
0 & 0 & 0 & 0 & 0 & 0 & 1 & 0 & 0 & 0 & -1 & 0 & 0 & 0 & 0 \\
0 & 0 & 0 & 0 & 0 & 0 & 0 & 1 & 0 & 0 & 0 & -1 & 0 & 0 & 0 \\
0 & 0 & 0 & 0 & 0 & 0 & 0 & 0 & 1 & 0 & 0 & 0 & -1 & 0 & 0 \\
0 & 0 & 0 & 0 & 0 & 0 & 0 & 0 & 0 & 1 & 0 & 0 & 0 & -1 & 0 \\
0 & 0 & 0 & 0 & 0 & 0 & 0 & 0 & 0 & 0 & 1 & 0 & 0 & 0 & -1 \\
1 & -1 & 0 & 0 & 0 & 0 & 0 & 0 & 0 & 0 & 0 & 0 & 0 & 0 & 0 \\
0 & 0 & 0 & 0 & 1 & -1 & 0 & 0 & 0 & 0 & 0 & 0 & 0 & 0 & 0 \\
0 & 0 & 0 & 0 & 0 & 0 & 0 & 0 & 1 & -1 & 0 & 0 & 0 & 0 & 0 \\
0 & 0 & 0 & 0 & 0 & 0 & 0 & 0 & 0 & 0 & 0 & 0 & 1 & -1 & 0 \\
0 & 1 & -1 & 0 & 0 & 0 & 0 & 0 & 0 & 0 & 0 & 0 & 0 & 0 & 0 \\
0 & 0 & 0 & 0 & 0 & 1 & -1 & 0 & 0 & 0 & 0 & 0 & 0 & 0 & 0 \\
0 & 0 & 0 & 0 & 0 & 0 & 0 & 0 & 0 & 1 & -1 & 0 & 0 & 0 & 0 \\
0 & 0 & 0 & 0 & 0 & 0 & 0 & 0 & 0 & 0 & 0 & 0 & 0 & 1 & -1 \\
0 & 0 & 1 & -1 & 0 & 0 & 0 & 0 & 0 & 0 & 0 & 0 & 0 & 0 & 0 \\
0 & 0 & 0 & 0 & 0 & 0 & 1 & -1 & 0 & 0 & 0 & 0 & 0 & 0 & 0 \\
0 & 0 & 0 & 0 & 0 & 0 & 0 & 0 & 0 & 0 & 1 & -1 & 0 & 0 & 0
\end{pmatrix}^T \begin{pmatrix}
c_g(t)\, N(t) \\
k_{00\to01}\, \tilde{x}_{g,00} \\
k_{00\to10}\, \tilde{x}_{g,00} \\
k_{01\to02}\, \tilde{x}_{g,01} \\
k_{01\to11}\, \tilde{x}_{g,01} \\
k_{02\to03}\, \tilde{x}_{g,02} \\
k_{02\to12}\, \tilde{x}_{g,02} \\
k_{03\to13}\, \tilde{x}_{g,03} \\
k_{10\to11}\, \tilde{x}_{g,10} \\
k_{10\to20}\, \tilde{x}_{g,10} \\
k_{11\to12}\, \tilde{x}_{g,11} \\
k_{11\to21}\, \tilde{x}_{g,11} \\
k_{12\to13}\, \tilde{x}_{g,12} \\
k_{12\to22}\, \tilde{x}_{g,12} \\
k_{13\to23}\, \tilde{x}_{g,13} \\
k_{20\to21}\, \tilde{x}_{g,20} \\
k_{20\to30}\, \tilde{x}_{g,20} \\
k_{21\to22}\, \tilde{x}_{g,21} \\
k_{21\to31}\, \tilde{x}_{g,21} \\
k_{22\to23}\, \tilde{x}_{g,22} \\
k_{22\to32}\, \tilde{x}_{g,22} \\
k_{30\to31}\, \tilde{x}_{g,30} \\
k_{31\to32}\, \tilde{x}_{g,31} \\
d_{K27,1}\, \tilde{x}_{g,10} \\
d_{K27,1}\, \tilde{x}_{g,11} \\
d_{K27,1}\, \tilde{x}_{g,12} \\
d_{K27,1}\, \tilde{x}_{g,13} \\
d_{K27,2}\, \tilde{x}_{g,20} \\
d_{K27,2}\, \tilde{x}_{g,21} \\
d_{K27,2}\, \tilde{x}_{g,22} \\
d_{K27,2}\, \tilde{x}_{g,23} \\
d_{K27,3}\, \tilde{x}_{g,30} \\
d_{K27,3}\, \tilde{x}_{g,31} \\
d_{K27,3}\, \tilde{x}_{g,32} \\
d_{K36,1}\, \tilde{x}_{g,01} \\
d_{K36,1}\, \tilde{x}_{g,11} \\
d_{K36,1}\, \tilde{x}_{g,12} \\
d_{K36,1}\, \tilde{x}_{g,13} \\
d_{K36,2}\, \tilde{x}_{g,02} \\
d_{K36,2}\, \tilde{x}_{g,12} \\
d_{K36,2}\, \tilde{x}_{g,22} \\
d_{K36,2}\, \tilde{x}_{g,32} \\
d_{K36,3}\, \tilde{x}_{g,03} \\
d_{K36,3}\, \tilde{x}_{g,13} \\
d_{K36,3}\, \tilde{x}_{g,23}
\end{pmatrix}.$$

The model comprises $n_\psi = 29$ parameters

$$\boldsymbol{\psi} = (c, d_{K27,1}, d_{K27,2}, d_{K27,3}, d_{K36,1}, d_{K36,2}, d_{K36,3}, k_{00\to01}, k_{00\to10}, k_{01\to02}, k_{01\to11},$$
$$k_{02\to03}, k_{02\to12}, k_{03\to13}, k_{10\to11}, k_{10\to20}, k_{11\to12}, k_{11\to21}, k_{12\to13}, k_{12\to22},$$
$$k_{13\to23}, k_{20\to21}, k_{20\to30}, k_{21\to22}, k_{21\to31}, k_{22\to23}, k_{22\to32}, k_{30\to31}, k_{31\to32}),$$

which are required for the simulation of the observable. However, the relative weights are not independent and, thus, the model comprises only 28 independent parameters.

To bring the system to relative scale, we divide the total abundance of modifications by the number of histone tails

$$x_{g,ij} = \frac{\tilde{x}_{g,ij}}{N},$$
$$\dot{x}_{g,ij} = \frac{\dot{\tilde{x}}_{g,ij}}{N} - \frac{\tilde{x}_{g,ij}\dot{N}}{N^2}.$$

This yields for the relative scale

$$\begin{aligned}
\dot{x}_{g,ij} = \;& \chi_{\{(i,j)=(0,0)\}}(i,j)\, c_g(t) - c_g(t)\frac{\tilde{x}_{g,ij}}{N} \\
&+ \chi_{\{i>0\wedge(i,j)\neq(3,3)\}}(i,j)\, k_{i-1\,j\to i\,j}\, \frac{\tilde{x}_{g,i-1\,j}}{N} \\
&+ \chi_{\{j>0\wedge(i,j)\neq(3,3)\}}(i,j)\, k_{ij-1\to ij}\, \frac{\tilde{x}_{g,ij-1}}{N} \\
&- \chi_{\{i<3\wedge(i,j)\neq(2,3)\}}(i,j)\, k_{ij\to i+1\,j}\, \frac{\tilde{x}_{g,ij}}{N} \\
&- \chi_{\{j<3\wedge(i,j)\neq(3,2)\}}(i,j)\, k_{ij\to ij+1}\, \frac{\tilde{x}_{g,ij}}{N} \\
&+ \chi_{\{i<3\wedge(i,j)\neq(2,3)\}}(i,j)\, d_{K27,i+1}\, \frac{\tilde{x}_{g,i+1\,j}}{N} \\
&+ \chi_{\{j<3\wedge(i,j)\neq(3,2)\}}(i,j)\, d_{K36,j+1}\, \frac{\tilde{x}_{g,ij+1}}{N} \\
&- \chi_{\{i>0\wedge(i,j)\neq(3,3)\}}(i,j)\, d_{K27,i}\, \frac{\tilde{x}_{g,ij}}{N} \\
&- \chi_{\{j>0\wedge(i,j)\neq(3,3)\}}(i,j)\, d_{K36,j}\, \frac{\tilde{x}_{g,ij}}{N} \\
= \;& \chi_{\{(i,j)=(0,0)\}}(i,j)\, c_g(t) - c_g(t)x_{g,ij} \\
&+ \chi_{\{i>0\wedge(i,j)\neq(3,3)\}}(i,j)\, k_{i-1\,j\to i\,j}\, x_{g,i-1\,j} \\
&+ \chi_{\{j>0\wedge(i,j)\neq(3,3)\}}(i,j)\, k_{ij-1\to ij}\, x_{g,ij-1} \\
&- \chi_{\{i<3\wedge(i,j)\neq(2,3)\}}(i,j)\, k_{ij\to i+1\,j}\, x_{g,ij} \\
&- \chi_{\{j<3\wedge(i,j)\neq(3,2)\}}(i,j)\, k_{ij\to ij+1}\, x_{g,ij} \\
&+ \chi_{\{i<3\wedge(i,j)\neq(2,3)\}}(i,j)\, d_{K27,i+1}\, x_{g,i+1\,j} \\
&+ \chi_{\{j<3\wedge(i,j)\neq(3,2)\}}(i,j)\, d_{K36,j+1}\, x_{g,ij+1} \\
&- \chi_{\{i>0\wedge(i,j)\neq(3,3)\}}(i,j)\, d_{K27,i}\, x_{g,ij} \\
&- \chi_{\{j>0\wedge(i,j)\neq(3,3)\}}(i,j)\, d_{K36,j}\, x_{g,ij},
\end{aligned}$$

for $g = 1, 2, 3$ and $i, j = 0, 1, 2, 3$. At relative scale, (Eq. 2) can be seen as the dilution that occurs due to cell division.

The observables, i.e., the measurable output of the model, are the methylation ratios obtained by

$$y_{g,ij} = \frac{x_{g,ij}}{\sum_{i,j} x_{g,ij}}.$$

We assumed that methylation rates not change for different culture media and different dynamics were obtained solely due to presence/absence of dilution (Eq. 2) and different initial conditions (Eq. 3).

#### 2 Domain model

As an alternative, we considered that demethylation of K27 and K36 does not exist. For this, we proposed a model which assumes that certain domains of the chromatin are determined to acquire certain methylation patterns, e.g., due to particular transcription factor binding or parental histone context. For example, histones of domain *00* do not get any methylations at all; histones of the domain *20* will get no additional methylations once the *20* state is reached; histones of the domain *31* can acquire the 31 methylation via different pathways (Fig. 2B). The histone composition tends towards the state where all domains acquired their determined state. Since newly incorporated histones are unmodified (*00*), the model still shows dynamics.

To construct this domain model, we denoted $w_{lm}$ as the relative size of domain $lm$, with $\sum_{l,m} w_{lm} = 1$. Let $x^{lm}_{g,ij}$ be the relative abundance of histones tails with methylation K27me$i$K36me$j$ in domain $lm$ for generation $g$. Then the ODEs are

$$\begin{aligned} \dot{x}^{lm}_{g,ij} = \; & \chi_{\{(i,j)=(0,0)\}}(i,j)\, c_g(t) - c_g(t)\, x^{lm}_{g,ij} \\ + \; & \chi_{\{0<i\leq l\}}(i,j)\, k_{i-1\,j\to ij}\, x^{lm}_{g,i-1\,j} \\ + \; & \chi_{\{0<j\leq m\}}(i,j)\, k_{ij-1\to ij}\, x^{lm}_{g,ij-1} \\ - \; & \chi_{\{i<l\}}(i,j)\, k_{ij\to i+1\,j}\, x^{lm}_{g,ij} \\ - \; & \chi_{\{j<m\}}(i,j)\, k_{ij\to ij+1}\, x^{lm}_{g,ij}\,, \end{aligned}$$

with $c_g(t)$ as defined in (Eq. 2) and initial conditions

$$x^{lm}_{g,ij}(t) = 0 \text{ for } \begin{cases} t < t_0 & g = 1\,, \\ t < -3\,\mathrm{h} & g = 2\,, \\ t < 0\,\mathrm{h} & g = 3\,, \end{cases} \qquad \forall i,j,l,m\,. \tag{4}$$

The observables are obtained by

$$y_{g,ij} = \frac{\sum_{l,m} w_{lm} x^{lm}_{g,ij}}{\sum_{l,m} w_{lm} \sum_{i,j} x^{lm}_{g,ij}}.$$

We assumed that the methylation rate constants are shared between the domains, and estimated them together with the relative sizes of the domains from the data. The model comprises $n_\psi = 38$ parameters

$$\begin{aligned} \boldsymbol{\psi} = (& c, k_{00\to01}, k_{00\to10}, k_{01\to02}, k_{01\to11}, k_{02\to03}, k_{02\to12}, k_{03\to13}, k_{10\to11}, k_{10\to20}, k_{11\to12}, k_{11\to21}, \\ & k_{12\to13}, k_{12\to22}, k_{13\to23}, k_{20\to21}, k_{20\to30}, k_{21\to22}, k_{21\to31}, k_{22\to23}, k_{22\to32}, k_{30\to31}, k_{31\to32}, \\ & w_{00}, w_{01}, w_{02}, w_{03}, w_{10}, w_{11}, w_{12}, w_{13}, w_{20}, w_{21}, w_{22}, w_{23}, w_{30}, w_{31}, w_{32})\,, \end{aligned}$$

which are required for the simulation of the observable.

#### 3 Model calibration

Experimental measurements are generally noise corrupted and the model needs to take this into account. It has been shown that often a Laplace measurement noise model outperforms a Gaussian noise model (Maier et al., 2017). Therefore, we first compared the Gaussian and Laplace distributed measurement noise model. We compared the model output and the observables on a log-scale and offsetted both to cope with zero measurements

$$\log(\bar{y}_{g,ij} + \text{offset}) \sim p\left(\log(\bar{y}_{g,ij} + \text{offset}) | \log\left(y_{g,ij} + \text{offset}\right), \sigma\right) , \tag{5}$$

with noise distribution $p$. The model parameters, including methylation rate constants, cell cycle and measurement noise, are comprised in the parameter vector $\boldsymbol{\theta}$. The measurement for the time point indexed by $k$ is denoted by $\bar{y}^k_{g,ij}$. The log-likelihood assuming Gaussian noise is given by

$$\log \mathcal{L}(\boldsymbol{\theta}) = \log \mathcal{L}_{\mathcal{D}}(\boldsymbol{\theta}) = -\frac{1}{2} \sum_{g,i,j,k} \left( \log\left(2\pi\sigma^2\right) + \left( \frac{\log(\bar{y}^k_{g,ij} + \text{offset}) - \log\left(y_{g,ij}(t_k, \boldsymbol{\theta}) + \text{offset}\right)}{\sigma} \right)^2 \right) , \tag{6}$$

and for Laplace noise by

$$\log \mathcal{L}(\boldsymbol{\theta}) = - \sum_{g,i,j,k} \left( \log\left(2\sigma\right) + \frac{|\log(\bar{y}^k_{g,ij} + \text{offset}) - \log\left(y_{g,ij}(t_k, \boldsymbol{\theta}) + \text{offset}\right)|}{\sigma} \right) . \tag{7}$$

The optimal parameters are obtained by maximizing the likelihood function, yielding the maximum likelihood estimate (MLE) $\hat{\boldsymbol{\theta}}$. Maximum likelhood estimation was performed using the parameter estimation toolbox PESTO (Stapor et al., 2018), which provides an interface to the MATLAB function `fmincon`. For numerical reasons, we transformed parameters which are supposed to be positive to a logarithmic scale (Hass et al., 2019). The models were simulated using AMICI (Fröhlich et al., 2017), which provides an interface to the SUNDIALS solver suite (Hindmarsh et al., 2005). For model comparison, we employed the Bayesian Information Criterion (BIC) (Schwarz, 1978). The BIC value for model indexed by $m$ is

$$\text{BIC}_m = -2 \log \mathcal{L}(\hat{\boldsymbol{\theta}}_m) + \log\left(n_{\mathcal{D}}\right) n_{\theta_m} , \tag{8}$$

with $n_{\mathcal{D}}$ data points. The BIC rewards good likelihood values and penalizes the number of parameters.

We chose offset $= 10^{-1}$ which provided the best fit with respect to the QQ-plots for the global model and the domain model with 15 domains using Laplace noise (Fig. S2A&B). We performed 100 local optimization runs for Gaussian and Laplace noise using the hierarchical approach for optimization proposed by Loos et al. (2018). For the hierarchical approach, we analytically calculated the measurement noise parameter $\sigma$, which is shared for all time points, observables and generations. The hierarchical approach outperformed the standard approach for optimization. We found strong support for Laplace noise over Gaussian noise for both models ($\Delta$BIC $> 600$). Both models were able to describe the data from untreated cells well (Fig. 2C and S2C&D). However, a model without demethylation for K27 was not able to describe, e.g., levels of K27me3K36me0 or K27me3K36m1 (Fig. S2C). Since only the domain model is able to explain inhibitor experiments (Fig. 2E), we reject the global model for the description of K27K36 dynamics.

#### 4 Model reduction and averaging

We did not expect all 15 domains to be necessary to explain the data and thus the domain model could be overparametrized. Since it was unclear which domains exist a priori, we performed model selection to detect the present domains. If we consider all potential combinations of domains, we would end up with $2^{15}$ models, which is computationally too expensive. Performing a combination of forward-selection and backward-elimination, we found eight domains $01, 02, 03, 13, 21, 23, 30, 32$ which were necessary to explain the data. To be robust against the precise choice of domains, we performed model averaging over all models which were calibrated in the process of model reduction. For this, the BIC weight,

$$\omega_s = \frac{\exp(-\frac{1}{2}\mathrm{BIC}_s)}{\sum_{\tilde{s}=1}^{n_M} \exp(-\frac{1}{2}\mathrm{BIC}_{\tilde{s}})} , \tag{9}$$

of each of the $n_M = 137$ models was employed (Fig. S2H). Since the rates are not comparable across models due to different domains, we analyzed the fluxes, which are the product of the abundance of the state and the methylation rate constant

$$\mathrm{flux}_{ij\to i+1j} = \sum_{l=0}^{i+1} \sum_{m=0}^{j} x_{1,ij}^{lm}(t_0) \cdot k_{ij\to i+1j} ,$$
$$\mathrm{flux}_{ij\to ij+1} = \sum_{l=0}^{i} \sum_{m=0}^{j+1} x_{1,ij}^{lm}(t_0) \cdot k_{ij\to i+1j} .$$

The fluxes shown in Fig. 2F model averaged using the BIC weights (Eq. 9) (Wassermann, 2000).

#### 5 Model prediction and validation

To further test and validate the models, we used the data of the inhibitor experiment. Using our calibrated models, we predicted the total amount of K27me3 in generation 1 under inhibitor treatment (Fig. 2D). For this, we assumed that the tri-methylation rates were inhibited by a factor $\kappa$:

$$k_{2j\to 3j,\mathrm{inh}} = (1-\kappa) \cdot k_{2j\to 3j,\mathrm{untr}} . \tag{10}$$

To obtain reasonable values for $\kappa$, we analyzed a simplified model which only considers K27 methylations of histones of generation 3, $x_0, x_1, x_2, x_3$ for un-, mono-, di-, and tri-methylation at K27 and assumed independence between the methylation sites. The model reads

$$\dot{x}_0 = c - cx_0 - k_{0\to 1}\, x_0 + d_{K27,1}x_1 ,$$
$$\dot{x}_1 = -cx_1 + k_{0\to 1}\, x_0 - k_{1\to 2}\, x_1 + d_{K27,2}x_2 - d_{K27,1}x_1 ,$$
$$\dot{x}_2 = -cx_2 + k_{1\to 2}\, x_1 - k_{2\to 3}\, x_2 + d_{K27,3}x_3 - d_{K27,2}x_2 ,$$
$$\dot{x}_3 = -cx_3 + k_{2\to 3}\, x_2 - d_{K27,3}x_3 .$$

The steady states for K27me3 for untreated cells and cells in the inhibitory experiment are given by

$$x_{3,\mathrm{untr}} = \frac{k_{2\to 3}\, x_{2,\mathrm{untr}}}{d_{K27,3} + c} , \tag{11}$$

$$x_{3,\mathrm{inh}} = \frac{(1-\kappa) k_{2\to 3}\, x_{2,\mathrm{inh}}}{d_{K27,3} + c} . \tag{12}$$

Thus, K27me3 only depends on the tri-methylation rate constant $k_{2\to 3}$, the demethylation rate constant $d_{K27,3}$, the dilution rate constant $c$, the amount of relative K27me2 and in the inhibitor case the factor $\kappa$ (Eq. 10) by which the tri-methylation is inhibited (Fig. S2E). Thus we obtained

$$\frac{d_{K27,3} + c}{k_{2\to 3}} = \frac{x_{2,\mathrm{untr}}}{x_{3,\mathrm{untr}}} = (1-\kappa)\frac{x_{2,\mathrm{inh}}}{x_{3,\mathrm{inh}}} \tag{13}$$

$$\Rightarrow \kappa = 1 - \frac{x_{2,\mathrm{untr}}\, x_{3,\mathrm{inh}}}{x_{3,\mathrm{untr}}\, x_{2,\mathrm{inh}}} . \tag{14}$$

Using the last time point for generation 3 from data $\mathcal{D}_{\text{inh}}$, we obtained for three replicates a rough estimate $\kappa = 0.928 \pm 0.052$ (Fig. S2G). The same expression for $\kappa$ (Eq. 14) is also valid for the domain model.

We compared our model predictions to the experimental data for K27me3 levels, i.e., summing all states with K27me3, to be robust against potential effects of the inhibitor on K36 methylations. We only assumed the tri-methylation rate to change. However, if also mono-, or di-methylation changes, the model predictions would be even lower and the illustrated predictions in Fig. 2E can be seen as rough estimates for the upper bound.

#### 6 Modeling differences between generation 2 histones in untreated and recovery experiments

For all further analyses, we used a domain model with eight domains $(01, 02, 03, 13, 21, 23, 30, 32)$. To detect differences between generation 2 histones of untreated and recovering cells, we performed a forward selection to find the most substantial changes in rate constants. In the first step of forward selection, only changes in the rate constants $k_{00\to10}$ and $k_{20\to30}$ yield substantial improvements with $\Delta$BIC$<$ 10. Allowing the rate constant for $k_{00\to10}$ to differ yields the biggest improvement in BIC values, and this rate constant is estimated to be roughly halved (0.47) in cells lacking H3K27me3. Overall, the model allowing for differences in $k_{00\to10}, k_{00\to01}, k_{20\to21}$ and $k_{20\to30}$ has the lowest BIC and thus the highest quality (Fig. 3E-F).

#### 7 Implementation

The MATLAB code used for the analysis of the manuscript will be made available via CodeOcean. The toolboxes used for the ODE simulation (AMICI) and parameter estimation (PESTO) are both available under `https://github.com/ICB-DCM`. The analysis was performed with MATLAB 2017b.
