## Supplemental Table 1 for "Domain model explains propagation dynamics and stability of K27 and K36 methylation landscapes"

| Mass [m/z] | Formula [M] | Formula type | Species | CS [z] | Polarity | Start [min] | End [min] | (N)CE | (N)CE type | MSX ID | Comment |
|---|---|---|---|---|---|---|---|---|---|---|---|
| 539,3141 | | | | 3 | Positive | 35 | 45 | 29 | NCE | | H3_27_40_K27K36K37_m1pp + H3_27_40_K27K36K37_pm1p_light |
| 545,346 | | | | 3 | Positive | 35 | 45 | 29 | NCE | | H3_27_40_K27K36K37_m1pp + H3_27_40_K27K36K37_pm1p_medium |
| 550,6644 | | | | 3 | Positive | 35 | 45 | 29 | NCE | | H3_27_40_K27K36K37_m1pp + H3_27_40_K27K36K37_pm1p_heavy |
| 529,9825 | | | | 3 | Positive | 30 | 40 | 30 | NCE | | H3_27_40_K27K36K37_m2m1p + H3_27_40_K27K36K37_pm3p_light |
| 536,0143 | | | | 3 | Positive | 30 | 40 | 30 | NCE | | H3_27_40_K27K36K37_m2m1p + H3_27_40_K27K36K37_pm3p_medium |
| 541,3328 | | | | 3 | Positive | 30 | 40 | 30 | NCE | | H3_27_40_K27K36K37_m2m1p + H3_27_40_K27K36K37_pm3p_heavy |
| 534,6544 | | | | 3 | Positive | 32 | 40 | 30 | NCE | | H3_27_40_K27S28K36K37_me3me1p + H3_27_40_K27S28K36K37_me1me3p_light |
| 540,6862 | | | | 3 | Positive | 32 | 40 | 30 | NCE | | H3_27_40_K27S28K36K37_me3me1p + H3_27_40_K27S28K36K37_me1me3p_medium |
| 546,0047 | | | | 3 | Positive | 32 | 40 | 30 | NCE | | H3_27_40_K27S28K36K37_me3me1p + H3_27_40_K27S28K36K37_me1me3p_heavy |
| 525,3106 | | | | 3 | Positive | 30 | 38 | 30 | NCE | | H3_27-40_K27K36K37_pme2p + H3_27-40_K27K36K37_me2pp_light |
| 531,3424 | | | | 3 | Positive | 30 | 38 | 30 | NCE | | H3_27-40_K27K36K37_pme2p + H3_27-40_K27K36K37_me2pp_medium |
| 536,6609 | | | | 3 | Positive | 30 | 38 | 30 | NCE | | H3_27-40_K27K36K37_pme2p + H3_27-40_K27K36K37_me2pp_heavy |
| 520,6509 | | | | 3 | Positive | 25 | 35 | 32 | NCE | | H3_27-40_K27K36K37_me2me3p + H3_27-40_K27K36K37_me3me2p_light |
| 526,6827 | | | | 3 | Positive | 25 | 35 | 32 | NCE | | H3_27-40_K27K36K37_me2me3p + H3_27-40_K27K36K37_me3me2p_medium |
| 532,0011 | | | | 3 | Positive | 25 | 35 | 32 | NCE | | H3_27-40_K27K36K37_me2me3p + H3_27-40_K27K36K37_me3me2p_heavy |
| 543,986 | | | | 3 | Positive | 34 | 42 | 30 | NCE | | H3_27_40_K27K36K37_m1m1p_light |
| 550,0178 | | | | 3 | Positive | 34 | 42 | 30 | NCE | | H3_27_40_K27K36K37_m1m1p_medium |
| 555,3363 | | | | 3 | Positive | 34 | 42 | 30 | NCE | | H3_27_40_K27K36K37_m1m1p_heavy |
